# Female Specific Restrictive Cardiomyopathy and Metabolic Dysregulation in transgenic mice expressing a Peptide of the Amino-Terminus of GRK2

**DOI:** 10.1101/2024.10.14.618348

**Authors:** Kamila M. Bledzka, Iyad H. Manaserh, Amanda D. Ifft, Julie H. Rennison, Madison Bohacek, Katie M. Vasiliauskas, Jessica Grondolsky, Isaac Ampong, David R. Van Wagoner, Sarah M. Schumacher

**Author notes:** **Corresponding Author:** Sarah M. Schumacher, PhD Assistant Staff, Cardiovascular & Metabolic Sciences Assistant Professor, Molecular Medicine Cleveland Clinic Lerner College of Medicine, CWRU Lerner Research Institute, NB50 Cleveland Clinic, 9500 Euclid Avenue, Cleveland, Ohio 44195.

## Abstract

Cardiovascular disease and heart failure are a major health challenge, with sex differences in pathophysiology and treatment responses critically influencing patient outcomes. G protein-coupled receptor (GPCR) kinase 2 (GRK2) is a pivotal regulator of cellular signaling whose elevation is a hallmark of heart failure progression. Its complex network of protein interactions impact a wide range of physiological and pathophysiological processes including cardiac function. In this study, we examined the effects of cardiac-restricted expression of an amino-terminal peptide of GRK2 (βARKnt) in female mice subjected to acute and chronic pressure overload. Our findings reveal that that βARKnt affects hypertrophy development and cardiac function differently in female mice than in males, leading to a transition to heart failure not observed in control females or βARKnt males. Notably, the βARKnt female mice exhibited baseline hypertrophy with distinct left atrial morphology, increased fibrosis, and immune cell infiltration compared to the controls, which progressed under chronic stress, indicating adverse cardiac remodeling. Furthermore, βARKnt female mice, unlike males, exhibit impaired tissue respiration following acute pressure overload and altered glucose sensitivity and insulin tolerance, highlighting significant remodeling of cardiac and systemic metabolism.

## 1. Introduction

G protein-coupled receptor (GPCR) kinase 2 (GRK2) is a pivotal regulator of cellular signaling, with a complex interactome that influences a wide range of physiological processes including cardiac function and a variety of systemic diseases. This kinase is primarily known for its role in phosphorylating GPCRs, thus facilitating receptor desensitization and internalization. However, emerging research shows GRK2s unique functional domains extend beyond GPCR regulation, encompassing robust protein interactions with various non-GPCR targets.[1, 2] These domains include the central catalytic domain and distinct regulatory segments at both the amino and carboxyl terminals, each contributing to GRK2s multifaceted roles in cellular signaling and regulation. Although GRK2s functionality is well established, knowledge of the underlying signaling mechanisms *in vivo* is lacking. Understanding the intricate relationships with GRK2s interactome is crucial for elucidating its mechanistic involvement in cardiac function and various pathophysiological conditions, revealing potential therapeutic interventions. One notable domain is the Regulator of G protein Signaling (RGS) homology domain (amino acids 51-173). We have previously demonstrated that this domain can interact selectively with Gαq *in vivo*, sequestering Gαq and arresting downstream signaling through Gq-coupled GPCRs, ultimately decreasing cardiac hypertrophy and blocking the development of heart failure.[3] In addition to RGS/Gαq, it was previously shown that the carboxyl terminus (βARKct) can interact with G_βγ_ and heat shock protein 90.[4–6] Initial studies on transgenic mice expressing a specific fragment of GRK2s RGS domain, referred to as βARKnt (residues 50-145), showed that this shorter peptide provoked baseline hypertrophy and increased β-adrenergic receptor (βAR) density without modifying downstream signaling via Gαs or Gαq.[7] This further supported the idea of domain specific interactions for GRK2.[7, 8]

To further investigate the variety of protein interactions encompassed by these GRK2 subdomains and their functional consequences *in vivo*, we regenerated transgenic mice with cardiomyocyte specific expression of βARKnt, including a carboxyl-terminal flag tag to distinguish it from endogenous GRK2. In our initial study, male βARKnt mice were subjected to left ventricular pressure overload via transverse aortic constriction (TAC).[9] These mice again demonstrated baseline hypertrophy, but exhibited proportional adaptive left ventricular hypertrophy during pressure overload and, unlike their non-transgenic littermate controls (NLCs), did not transition to heart failure. Unlike GRK2 upregulation that is bad for the heart and inhibits insulin signaling, but causes an overall lean phenotype, βARKnt males exhibited a more physiological hypertrophy, with reduced abdominal fat weight and increased insulin signaling. These findings suggest that βARKnt has unique effects on cardiac hypertrophy and dysfunction under pressure overload, indicating potential novel interactions that contribute to cardioprotection during heart failure in males.[9]

Cardiovascular disease and heart failure present a significant health burden, yet sex differences in the pathophysiology and treatment responses are increasingly recognized as critical factors influencing patient outcomes. Despite these differences, women have historically been underrepresented in clinical trials, 20-30% enrollment over the past 4 decades, leading to a gap in understanding the unique biological and hormonal influences that may affect disease progression and treatment efficacy.[10] As the medical field begins to acknowledge the significance of sex-based differences in heart failure, there is an urgent call to incorporate more female subjects in research to illuminate the distinct pathophysiological mechanisms and signaling pathways involved in heart failure among women.[11] Addressing this disparity is essential for developing more effective, gender-personalized approaches to diagnosis and treatment, ultimately improving outcomes for all heart failure patients.

In this study, we subjected TgβARKnt female mice to both acute and chronic TAC to explore potential sex differences in the interactions of this peptide and its resulting effects on cardiac remodeling, with the goal of exposing novel therapeutic targets for heart failure. Our findings reveal that βARKnt influences hypertrophy and cardiac function differently in female mice than in males, resulting in a transition to heart failure not observed in control females.

## 2. Results

### 2.1 βARKnt transgenic female mice exhibit an elevated fractional shortening and baseline hypertrophy with a normal adaptive response to acute TAC

To generate the TgβARKnt line, the cDNA that encodes bovine GRK2 residues 50-145 was cloned into a vector driven by the αMHC promoter with a carboxyl-terminal flag-tag. Founder lines were established, and no gross phenotypic changes were observed in transgenic mice compared to non-transgenic littermate control (NLC) mice. Transgene expression was confirmed by Western blotting of cardiac lysates using a rabbit polyclonal flag antibody to detect the ∼16 kDa band (Figure 1A). To determine whether expression of this peptide of GRK2 would alter cardiac remodeling in an animal model of pressure overload, TgβARKnt and non-transgenic mice underwent TAC or Sham surgery. Echocardiography was performed at baseline, 2, and 4 weeks to monitor cardiac function and dimensions. Sufficient left ventricular (LV) pressure gradients after TAC were confirmed by pulsed-wave Doppler of the aortic arch 1-week post-surgery, with no differences between non-transgenic and Tg groups (Figure S1A). Cardiac function was assessed by measures of LV fractional shortening that were significantly elevated in βARKnt Sham mice at 2 and 4 weeks, and βARKnt TAC mice at baseline, 2, and 4 weeks, compared to their respective NLC controls (Figure 1B). Interestingly, while LV posterior wall thickness during diastole and systole was significantly higher in βARKnt mice prior to surgery, these measures were equivalently elevated in non-transgenic and Tg mice at 2 and 4 weeks after TAC compared to their respective Sham controls (Figure 1C, D). Accordingly, heart weight was greater in the βARKnt Sham and TAC mice 4 weeks after surgery (Figure 1E); however, when normalized to their respective Sham animals, the % increase in heart weight after TAC was the same (Figure 1F). Left atrial weight was also greater in the βARKnt Sham and TAC mice 4 weeks after surgery, but in contrast to heart weight, showed a significantly greater elevation in the Tg mice after TAC than non-transgenic controls (Figure 1G). No difference was observed in lung weight (Figure 1H) or tibia length (Figure S1B) across these groups. Together, these data demonstrate proportional LV hypertrophic growth in the βARKnt transgenic and non-transgenic mice in response to acute cardiac stress, with differential effects on left atrial remodeling.

**Figure 1:**
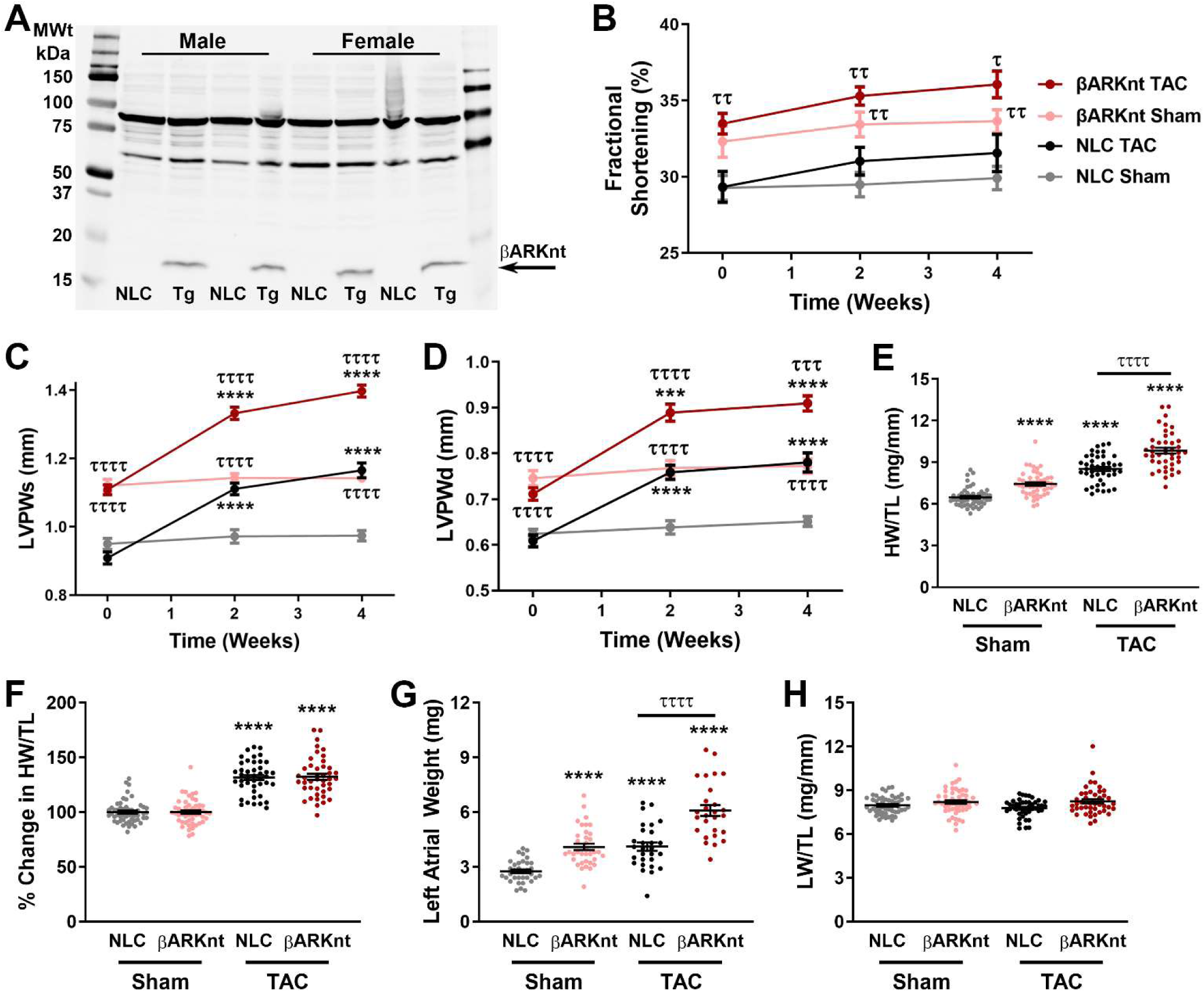
Cardiomyocyte-specific βARKnt expression enhances cardiac fractional shortening and elicits baseline hypertrophy with adaptive pressure overload-induced hypertrophy. **(A)** Western blot of male and female non-transgenic littermate control (NLC) and TgβARKnt ventricular lysates probed for flag-tagged βARKnt. **(B)** % left ventricular (LV) fractional shortening from NLC and βARKnt Sham and post-TAC animals at baseline, 2- and 4-weeks post-surgery. Serial measures of noted experimental groups for **(C)** LV posterior wall thickness during systole (LVPWs), and **(D)** diastole (LVPWd). ***, p < 0.001; ****, p < 0.0001 by two-way ANOVA with repeated measures and Tukey post-hoc test relative to corresponding Sham. ^τ^, p < 0.05; ^ττ^, p < 0.01; ^τττ^, p < 0.001; ^ττττ^, p < 0.0001 by two-way ANOVA with repeated measures and Tukey post-hoc test relative to corresponding NLC. n = 13-28 mice per group. Measures of **(E)** heart weight normalized to tibia length (HW/TL), **(F)** % change in HW/TL, **(G)** left atrial weight, and **(H)** lung weight normalized to TL (LW/TL) in these animals. ****, p < 0.0001 by one-way ANOVA with Tukey post-hoc test relative to NLC Sham. ^ττττ^, p < 0.0001 by one-way ANOVA with Tukey post-hoc test relative to corresponding NLC TAC. n = 26-58 mice per group.

### 2.2 βARKnt expression protects against decompensated hypertrophy and dilated cardiomyopathy, but not heart failure, after chronic TAC

To investigate whether the larger heart at baseline and after TAC induced by βARKnt expression would hasten the transition to heart failure, we followed a cohort of mice out to 14 weeks post-TAC. Cardiac structure and function were monitored by serial echocardiography every 2 weeks, and systolic pressure gradients were no different between the TAC groups 1-week post-surgery (Figure 2A). In contrast to non-transgenic male mice that transition to heart failure from 10-14 weeks after TAC,[9] cardiac fractional shortening (Figure 2B) was indistinguishable between non-transgenic Sham and TAC females across the 14-week time course. βARKnt Sham females exhibited a significantly higher fractional shortening than control, but these levels were consistent during the 14 weeks. Interestingly, despite a slight drop from 4 to 6 weeks, fractional shortening remained stable out to 14 weeks in the βARKnt TAC females as well. Further, while LV mass increased at 2 and 4 weeks then remained consistent in non-transgenic female mice after TAC, βARKnt mice exhibited a robust and continual increase across the 14-week time course (Figure 2C). This rapid departure in LV mass was not explained by LV posterior wall hypertrophy as measured during systole (LVPWs, Supplemental Figure 2A) or diastole (LVPWd, Supplemental Figure 2B), though, as the expansion of these measures was proportional between non-transgenic and βARKnt mice over time. Looking instead at measures of LV anterior wall thickness during systole (LVAWs, Supplemental Figure 2C) and diastole (LVAWd, Supplemental Figure 2D), there appeared to be a more significant increase in these measures in the βARKnt mice during chronic pressure overload.

**Figure 2:**
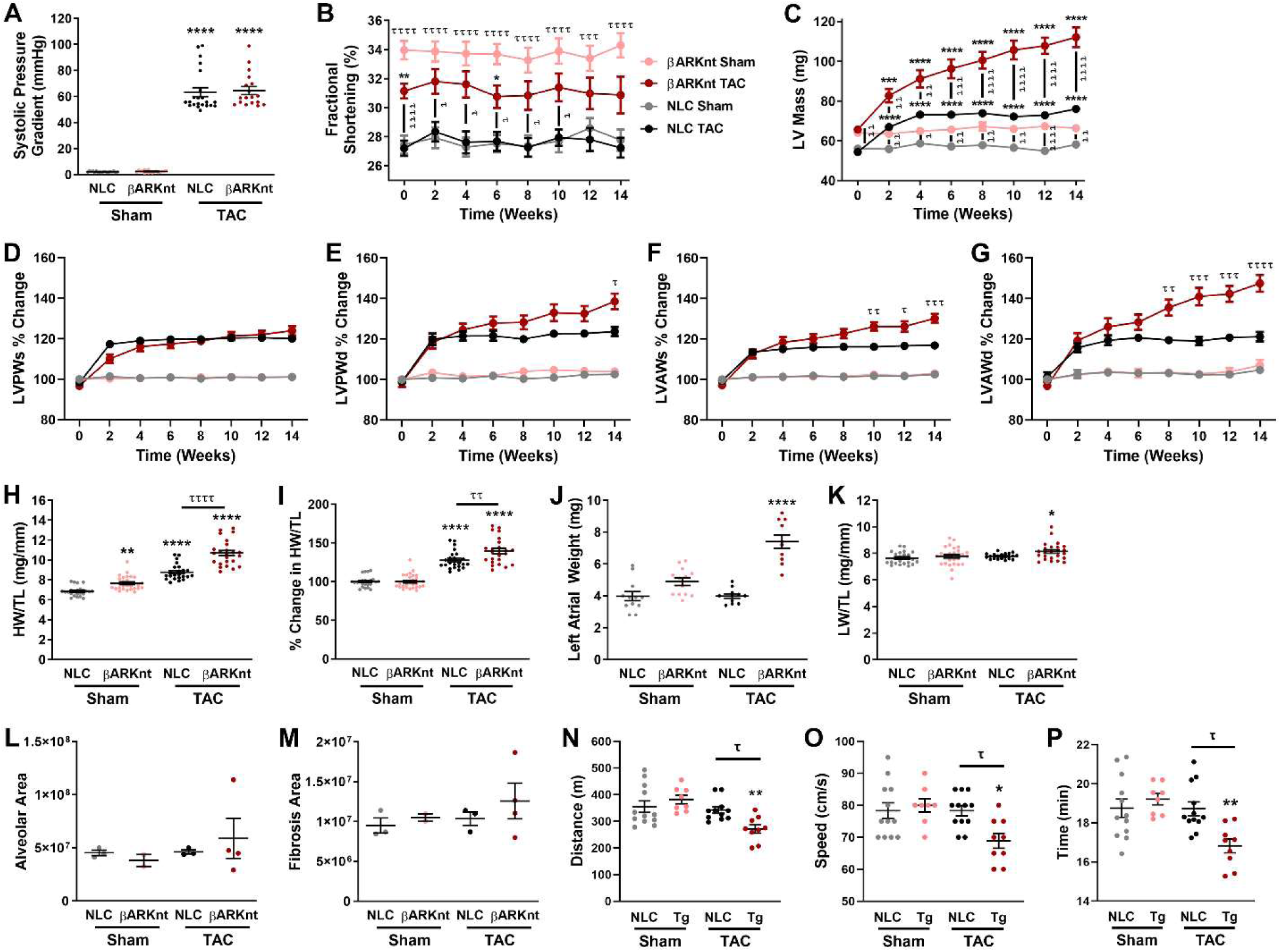
Despite preserved cardiac function, βARKnt hearts exhibit asymmetric hypertrophic cardiomyopathy and heart failure after chronic pressure overload. **(A)** Measures of the systolic pressure gradient in the aorta of non-transgenic littermate control (NLC) and TgβARKnt Sham and 14 weeks after TAC animals. Serial measures of noted experimental groups for **(B)** % left ventricular (LV) fractional shortening, **(C)** LV mass, and percentage increase over respective Sham in **(D)** LV posterior wall thickness during systole (LVPWs), **(E)** LV posterior wall thickness during diastole (LVPWd), **(F)** LV anterior wall thickness during systole (LVAWs), and **(G)** diastole (LVAWd). ***, p < 0.001; ****, p < 0.0001 by two-way ANOVA with repeated measures and Tukey post-hoc test relative to corresponding Sham. ^τ^, p < 0.05; ^ττ^, p < 0.01; ^τττ^, p < 0.001; ^ττττ^, p < 0.0001 by two-way ANOVA with repeated measures and Tukey post-hoc test relative to corresponding NLC. n = 16-20 mice per group. Measures of **(H)** heart weight normalized to tibia length (HW/TL), **(I)** % change in HW/TL, **(J)** left atrial weight, **(K)** lung weight normalized to tibia length (LW/TL), **(L)** alveolar area, **(M)** fibrotic area, **(N)** distance run, **(O)** running speed, and **(P)** running time in these animals. *, p < 0.05; **, p < 0.01; ***, p < 0.01; ****, p < 0.0001 by one-way ANOVA with Tukey post-hoc test relative to NLC Sham. ^τ^, p < 0.05; ^ττ^, p < 0.01; ^ττττ^, p < 0.0001 by one-way ANOVA with Tukey post-hoc test relative to NLC TAC. n = 7-29 mice per group.

To gain more clarity on the parameters contributing to the substantial divergence in LV mass, we normalized all these wall thickness measures to their own sham controls and looked at when and where the percentage increase above control significantly deviated between non-transgenic and βARKnt mice after TAC. Interestingly, we observed no significant difference in LVPWs over the 14-week time course (Figure 2D), with a mild deviation in LVPWd beginning at 6 weeks that was only significant at 14 weeks (Figure 2E). LVAWs was divergent by 8 weeks at significant at 10, 12, and 14 (Figure 2F); however, the greatest departure was in LVAWd that was significant from 8 weeks on (Figure 2G). Together, these data demonstrate that hypertrophic growth in response to chronic pressure overload is greater in the anterior than posterior wall in the βARKnt female mice, and more notable as increased diastolic wall thickness. This contrasts with non-transgenic female mice that have equivalent and stable hypertrophic growth in both walls during contraction and relaxation. This is also in contrast to the stable, symmetric hypertrophy and cardioprotection observed in βARKnt male mice, and opposite to the switch from hypertrophy to wall thinning observed in non-transgenic male mice transitioning into maladaptive remodeling and dilated cardiomyopathy. However, the fact that this divergence in LV mass and wall thickness occurs from 8-14 weeks after TAC does suggest a transition in the remodeling process in the βARKnt, not control, females.

Heart weight normalized to tibia length was progressively increased in the βARKnt Sham and non-transgenic TAC mice with a further significant increase to βARKnt TAC (Figure 2H). Once normalized to their own Sham controls, the increase in heart weight was disproportionally greater in the βARKnt than non-transgenic mice, consistent with LV mass (Figure 2I). Despite a trend in the βARKnt Shams, left atrial weight was significantly increased only in the βARKnt TAC mice (Figure 2J). Further, lung weight normalized to tibia length was only elevated in the βARKnt TAC mice (Figure 2K). Together these data show a disproportional increase predominantly in anterior and diastolic wall thickness, and thus LV mass, in the βARKnt mice after chronic pressure overload. This is independent of any alteration in fractional shortening but was accompanied by a more pronounced increase in the normalized heart weight, atrial weight, and pulmonary congestion.

Due to the disproportional septal and diastolic hypertrophy and pulmonary congestion observed in the βARKnt female mice during chronic pressure overload, we evaluated pulmonary tissue health. Lungs were taken for histological analysis at 14 weeks after Sham or TAC surgery and subjected to Masson Trichrome staining. Quantification revealed no difference in left lobe alveolar area between control and βARKnt Sham or TAC mice (Figure 2L), but a trend towards an increase in fibrotic area in βARKnt TAC mice (Figure 2M). In addition, given the enhanced fractional shortening and yet increased ventricular remodeling, atrial weight, and pulmonary congestion suggestive of heart failure, we evaluated exercise tolerance in these mice. Treadmill exercise studies revealed no difference between Sham groups, but a significant decrease in distance run in both the control and βARKnt TAC mice that was more pronounced in the βARKnt mice (Figure 2N). Further, running speed (Figure 2O) and running time (Figure 2P) were only significantly reduced in the βARKnt TAC mice compared to both Sham and TAC controls, suggesting exercise intolerance in the βARKnt TAC females. Together, these data suggest that despite the preserved fractional shortening in the βARKnt mice during chronic pressure overload, they are still undergoing a transition to heart failure.

### 2.3 βARKnt promotes adverse ventricular remodeling including enhanced fibrosis and immune cell infiltration after acute and chronic TAC

To elucidate the underlying involvement of cardiomyocyte versus myocardial remodeling in the enhanced wall thickness and heart weight in the βARKnt mice, tissues were taken for histological and biochemical analysis 4- and 14-weeks post-TAC. To investigate the integrity of the myocardium, we performed Masson Trichrome staining of four chamber paraffin sections taken at the level of the aortic outflow tract. Representative images depict the greater baseline hypertrophy in the βARKnt mice, with a proportional increase in heart size and wall thickness 4 weeks after TAC, but a greater increase in wall thickness and atrial size 14 weeks after TAC (Figure 3A). Quantification of all images revealed a significant increase in LV myocyte area in the non-transgenic and βARKnt mice at 4 and 14 weeks after TAC (Figure 3B). Interestingly, LV fibrotic area was increased only in βARKnt Sham and TAC mice, not control, to an equivalent degree at 4 weeks and greater extent in the TAC mice at 14 weeks (Figure 3C). Thus, while total LV area was similarly elevated in the non-transgenic and βARKnt mice at 4 weeks, this increase was greater in the βARKnt than control mice at 14 weeks after TAC (Figure S3A). We noted distinct results when measuring these values in the left atrium of these hearts. Left atrial myocyte (Figure 3D), fibrotic (Figure 3E), and total (Figure S3B) areas trended towards an increase in non-transgenic TAC mice, with an equivalent (4 weeks) or greater (14 weeks) trend in βARKnt Sham animals, while only βARKnt TAC mice exhibited a significant increase in all 3 measures at both 4 and 14 weeks after TAC. In contrast, no difference was observed in right atrial myocyte (Figure S3C), fibrotic (Figure S3D), or total (Figure S3E) area between groups at either time point. Together, these data demonstrate that despite similar increases in myocyte area in the non-transgenic and βARKnt mice after TAC, interstitial LV fibrosis only comprises a sizable portion of the myocardium of βARKnt mice, and this proportion was only elevated after chronic pressure overload stress. Consistent with left atrial weight measures, these data also shown a trend (Sham) and significant (TAC) increase in left atrial myocyte and fibrotic areas where fibrotic area is greater at 14 weeks than 4 weeks.

**Figure 3:**
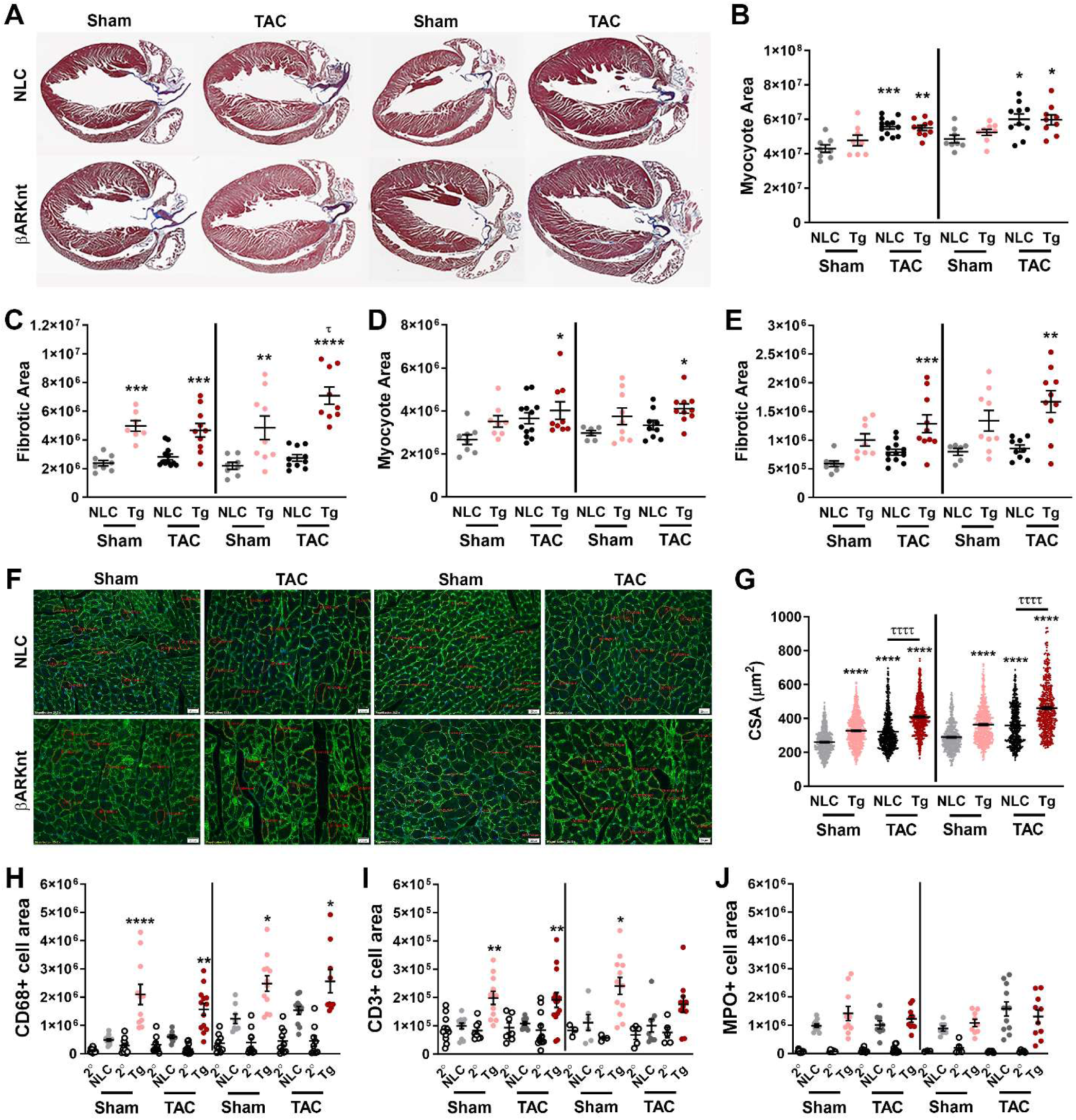
βARKnt hearts show elevated left ventricular and left atrial myocyte and fibrotic area at 4- and 14-weeks after TAC, with equivalently elevated myocyte cross-sectional area but differences in immune cell infiltration. **(A)** Representative images of Masson trichrome-stained murine heart sections in non-transgenic littermate control (NLC) and TgβARKnt Sham and TAC mice 4 (left) and 14 (right) weeks after surgery. Quantification of **(B)** myocyte area and **(C)** fibrotic area in the ventricles of these mice. Quantification of **(D)** myocyte area and **(E)** fibrotic area in the left atria of these mice. *, p < 0.05; **, p < 0.01; ***, p < 0.01; ****, p < 0.0001 by one-way ANOVA with Tukey post-hoc test relative to time-matched NLC Sham. ^τ^, p < 0.05 by one-way ANOVA with Tukey post-hoc test relative to time-matched βARKnt Sham. n = 9-15 hearts per group. **(F)** Representative images of WGA and 4′,6-diamidino-2-phenylindole (DAPI)-stained murine heart sections from NLC and βARKnt Sham and TAC mice 4 (left) and 14 (right) weeks after surgery. Scale bar, 20 µm. **(G)** Quantification of cardiomyocyte cross-sectional area (CSA) in these animals. ****, p < 0.0001 by one-way ANOVA with Tukey post-hoc test relative to time-matched NLC Sham. ^ττττ^, p < 0.0001 by one-way ANOVA with Tukey post-hoc test relative to time-matched NLC TAC. n = 489-722 cardiomyocytes from 12-15 hearts per group (∼45 myocytes per heart). Quantification of **(H)** CD68, **(I)** CD3, and **(J)** MPO positive cell area in these mice. *, p < 0.05; **, p < 0.01; ****, p < 0.0001 by one-way ANOVA with Tukey post-hoc test relative to time-matched NLC Sham. n = 3-12 mice per group.

Gene expression was measured via RT-PCR of cardiac mRNA. Atrial natriuretic peptide (ANP) (Figure S3F) and beta myosin heavy chain (βMHC) (Figure S3G) mRNA expression were significantly induced in βARKnt Sham and TAC hearts at 4 weeks and to a much greater extent 14 weeks after TAC, with a modest increase in control females at 14 weeks. There was no significant difference in B-cell lymphoma extra-large (Bcl-xL) (Figure S3H) or C/EBP homologous protein (CHOP) (Figure S3I) mRNA levels that may have indicated a shift towards an anti- or pro-apoptotic signaling, respectively. Transforming growth factor beta (TGFβ) was elevated in βARKnt Sham hearts only following chronic pressure overload (Figure S3J), whereas tumor necrosis factor alpha (TNFα) mRNA was similarly up in 4-week Sham and 14-week Sham and TAC βARKnt hearts (Figure S3K). Overall, these histological and mRNA data agree with our other tissue measures that some degree of cardiac hypertrophy and, unlike βARKnt male mice, fibrosis are evident even at baseline and more pronounced following chronic pressure overload stress.

To visualize the effect of βARKnt peptide expression on cardiomyocyte growth, we performed wheat germ agglutinin staining to measure myocyte cross-sectional area in these mice. These data revealed a significant increase in myocyte size in transgenic Sham animals at 4 weeks, demonstrating an alteration in cardiomyocyte size prior to cardiac stress (Figure 3F, G). These data also showed an even greater, but equivalent, increase in non-transgenic and βARKnt mice at 4 weeks and 14 weeks post-TAC. These data suggest that sources outside of cardiomyocyte hypertrophy, including fibrosis, are likely responsible for the disproportional increase in heart weight and total LV area observed in the βARKnt mice 14 weeks after TAC. Interestingly, these images revealed an increased abundance of small cells intermingled with the larger diameter cardiomyocytes. To determine the nature of these cells and whether they were infiltrating immune cells, we performed immunohistochemical staining for cluster of differentiation 68 (CD68), cluster of differentiation 3 (CD3), and myeloperoxidase (MPO). CD68 is a glycoprotein that is expressed on the surface of macrophages and is commonly used as a marker for macrophage activation, with increased CD68-positive macrophages associated with adverse cardiac remodeling and diastolic dysfunction. CD3 is a marker utilized for the identification of T cells, while myeloperoxidase (MPO) detects neutrophils. Although monocytes also express MPO, their expression levels are markedly lower compared to neutrophils.[12] Importantly, in contrast to males where CD68 positive staining was only elevated after TAC and to a similar extent in βARKnt and controls,[9] the βARKnt female mice demonstrated increased CD68 positive macrophage infiltration into the hearts before and after TAC and this was only significant in the βARKnt hearts (Figure 3H). CD3 positive T cell infiltration was elevated in both βARKnt Sham and TAC hearts after acute pressure overload, but this increased waned in the βARKnt TAC hearts during chronic stress (Figure 3I). The levels of MPO positive neutrophils were equivalent across all groups during acute and chronic pressure overload, demonstrating no significant genotype or stress response (Figure 3J). Together, these data suggest increase immune cell infiltration into the βARKnt hearts prior to and following cardiac stress that likely play a role in the remodeling phenotype.

Given the fibrosis, hypertrophy, and immune cell infiltration evident in 4-week Sham βARKnt hearts we decided to interrogate a cohort of younger mice. Hearts were harvested from 8-week-old male and female control and βARKnt mice. Heart weight normalized to tibia length revealed a trend, but not significant, increase in βARKnt male and female heart weight compared to control (Figure S4A), with no difference in tibia lengths at this age (Figure S4B). Interestingly, Masson Trichrome staining revealed that despite no difference in myocyte area (Figure S4C) βARKnt females, not males, already had significant myocardial fibrosis at 8 weeks of age (Figure S4D). At this age it was confined to the ventricles and not yet affecting the atria (Figure S4E, F). Immunohistochemistry also showed a significant increase in CD68 positive macrophage cell area in the βARKnt females only (Figure S4G), with no significant difference in the CD3 positive T cell area by sex or genotype (Figure S4H). Together, these data demonstrate that some degree of fibrosis and macrophage infiltration is present at an early age in the βARKnt female ventricular myocardium and thus contributes to the progressive remodeling after acute and chronic stress.

### 2.4 βARKnt female, not male, mice exhibit impaired tissue respiration after acute TAC and impaired glucose sensitivity and insulin tolerance after both acute and more so chronic TAC

Mitochondrial oxidative phosphorylation (OXPHOS) serves to generate ATP synthesis from the coupling of electron transport across the electron transport chain (ETC) complexes to phosphorylation of ADP by complex V (ATP synthetase). We utilized the standard substrate-uncoupler-inhibitor titration (SUIT) protocol in permeabilized myocardial free wall tissue sections from male and female Sham and TAC mice to quantify alterations in mitochondrial oxygen consumption.[13–15] Oxygen consumption in response to complex I substrates (malate, pyruvate, and glutamate) was significantly lower in βARKnt TAC females only, compared with control, sham operated and male hearts (Figure 4A). In response to malate and pyruvate alone, prior to glutamate, oxygen consumptions was significantly reduced equivalently in βARKnt Sham and NLC TAC females, with a more robust decrease in βARKnt TAC females, with no overall differences observed in males (Figure S5A). The subsequent responses to the complex II substrate succinate and the titration of the uncoupler FCCP (carbonyl cyanide p-trifluoromethoxyphenylhydrazone) that indicates maximum respiratory capacity were also reduced in the βARKnt TAC females only (Figure 4B, C). Following FCCP titration, respiration was inhibited with a combination of rotenone (inhibits complex I) and antimycin A (inhibits complex III). Rotenone and antimycin A were added to inhibit mitochondrial respiration, and non-mitochondrial respiration was subtracted from all other respiratory rates. Addition of the complex I inhibitor rotenone demonstrated a significant decrease in oxygen consumption by complex II also in βARKnt TAC females (Figure S5B). Subsequent complex III inhibitor antimycin A showed no overall differences between male and female groups under any conditions (Figure S5C). To measure complex IV-dependent respiration, first complex III-dependent respiration was inhibited by adding antimycin A (mitochondrial complex III inhibitor). Afterwards, complex IV-dependent respiration was stimulated by administering ascorbate and tetramethylphenylendiamine (TMPD). TMPD can auto-oxidize in the respiration buffer, therefore the maximal complex IV-dependent respiration rate was calculated by subtracting respiration rates before and after the addition of sodium azide, an inhibitor of mitochondrial complex IV. The addition of TMPD (a Cytochrome C electron donor) and ascorbate showed a trend, but not significant, decrease in βARKnt Sham and more so TAC females compared to control. Interestingly, this was significantly reduced compared to βARKnt TAC males under these conditions (Figure S5D). TMPD and ascorbate were quickly followed by azide to reveal azide-sensitive oxidation of TMPD by complex IV, with nearly identical results (Figure 4D).

**Figure 4:**
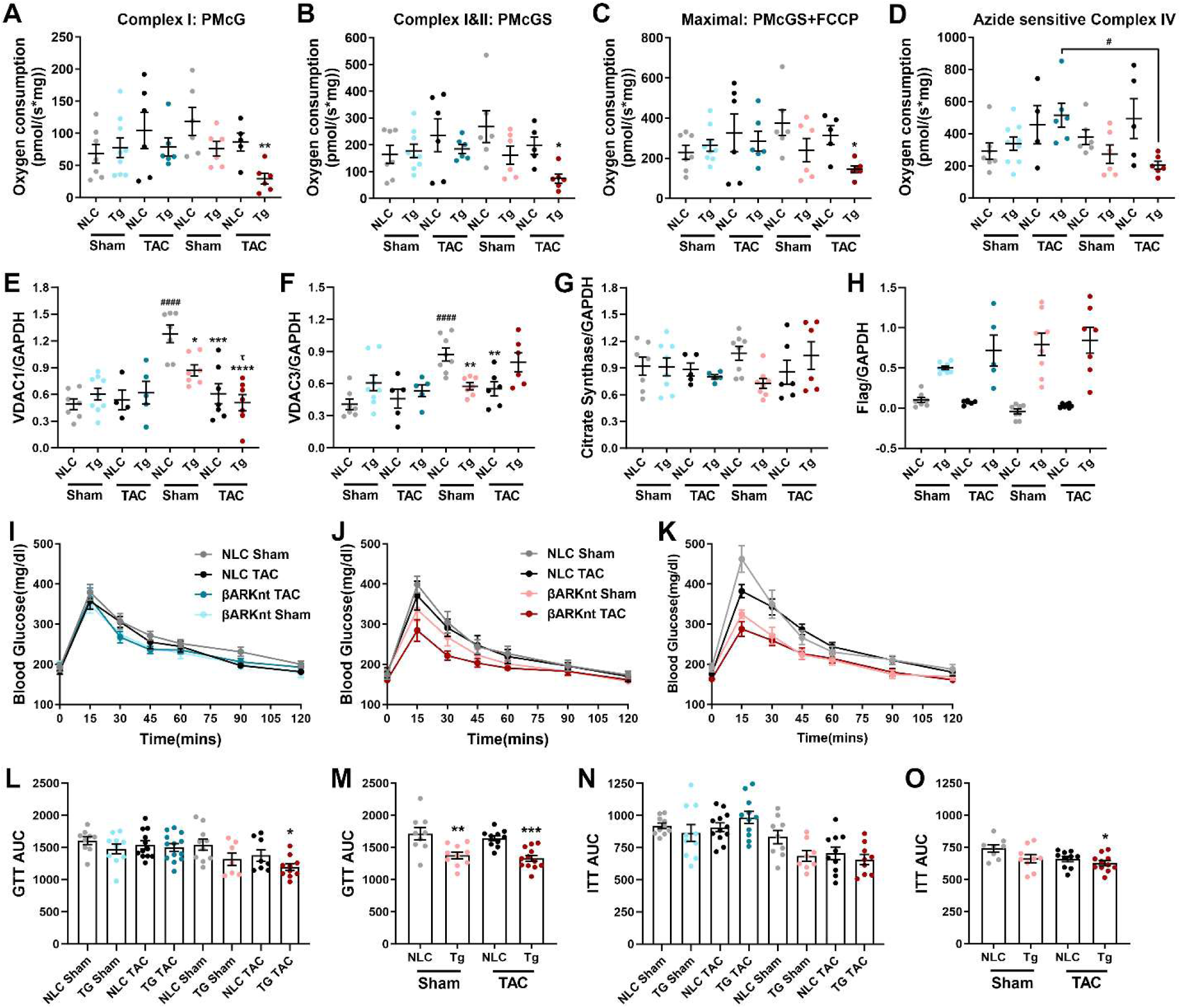
Sex and genotype differences in mitochondrial complex function, resident mitochondrial protein expression, and glucose and insulin handling at 4 and 14 weeks after surgery. Mitochondrial oxygen consumption in permeabilized cardiac muscle fibers from male and female non-transgenic littermate control (NLC) and TgβARKnt Sham and TAC mice 4 weeks after surgery in response to substrates and inhibitors of components of the electron transport chain (ETC) including **(A)** pyruvate, malate, ADP and glutamate (PMcG), **(B)** plus succinate (PMcGS), **(C)** plus FCCP, and **(D)** the difference between ascorbate + TMPD and subsequent sodium azide. n = 4-9 hearts per group. Quantification of **(E)** VDAC1, **(F)** VDAC3, **(G)** citrate synthase, and **(H)** Flag in male and female NLC and βARKnt ventricular tissue 4 weeks after surgery. All data were normalized to GAPDH within their respective blot. n = 4-9 per group from 4 Western blots. *, p < 0.05; **, p < 0.01; ***, p < 0.01; ****, p < 0.0001 by one-way ANOVA with Tukey post-hoc test relative to sex-matched NLC Sham. ^τ^, p < 0.05 by one-way ANOVA with Tukey post-hoc test relative to sex-matched βARKnt Sham. ^#^, p < 0.05; ^####^, p < 0.0001 by one-way ANOVA with Tukey post-hoc test relative to condition-matched male. Glucose tolerance test (GTT) measurements of blood glucose levels after IP injection of 2g/kg per BW of a sterile dextrose solution in **(I)** male and **(J)** female NLC and βARKnt Sham and TAC mice 4 weeks or **(K)** 14 weeks after surgery. Area under the curve (AUC) measurements of GTT in **(L)** male (left, blue) and female (right, pink) mice 4 weeks after TAC and **(M)** female mice 14 weeks after TAC. Area under the curve (AUC) measurements of ITT in **(N)** male (left, blue) and female (right, pink) mice 4 weeks after TAC and **(O)** female mice 14 weeks after TAC. *, p < 0.05; **, p < 0.01; ***, p < 0.01 by one-way ANOVA with Tukey post-hoc test relative to time- and sex-matched NLC Sham. n = 8-13 per group.

Summarized line graphs of these titration data show no major differences between males other than a mild trend towards an increase in oxygen consumption in both TAC groups compared to Sham (Figure S5E), compared to a mild downward trend in NLC TAC and βARKnt Sham females compared to NLC Sham and the significant depression in oxygen consumption in βARKnt TAC females under most conditions (Figure S5F). Of note, these graphs demonstrate that the NLC Sham females have the highest oxygen consumption whereas the NLC males have the lowest compared to other groups within their sex. This highlights the trend observed in all the individual titrations wherein the NLC Sham females showed a higher range of oxygen consumptions values than the corresponding NLC Sham males, suggesting baseline sex differences in mitochondrial respiration; however, this difference was lost after TAC. Together, these data demonstrate a reduction in specific components of the mitochondrial ETC (namely complexes I, II, and IV) in the βARKnt TAC females.

To investigate whether this decrease in mitochondrial OXPHOS was due more to decreased ETC complex function or potentially decreased mitochondrial content we performed RT-PCR of mitochondrial DNA markers ND1 and 16S relative to cellular 18S. These data revealed no significant difference in ND1 in NLC and βARKnt males 4 weeks after Sham or TAC surgery, but a significant increase in βARKnt TAC females at 4 weeks and decrease in NLC and much more so βARKnt females at 14 weeks after TAC (Figure S5G). Mitochondrial 16S was also no different in males but was significantly reduced in βARKnt Sham females at 4 weeks and equivalently in βARKnt Sham and TAC females 14 weeks after cardiac stress (Figure S5H). These data suggest that mitochondrial content does not differ dramatically by sex, but that cardiomyocyte βARKnt expression leads to a significant decrease in mitochondrial DNA that was differentially affected by TAC during acute and chronic stress. Additionally, we evaluated the protein levels of several resident mitochondrial proteins via Wester blotting. Voltage-dependent anion selective channel 1 (VDAC1), an outer mitochondrial and cell membrane ion channel, exhibited significantly more robust expression in control Sham females compared to males, but these levels were progressively reduced in βARKnt Sham, NLC TAC, and βARKnt TAC females at 4 weeks after surgery (Figure 4E). VDAC3 expression was also significantly greater in NLC Sham females compared to males but was only reduced in βARKnt Sham and NLC TAC females (Figure 4F). In contrast, the citric acid cycle pace-making enzyme citrate synthase was similarly expressed across all male and female groups (Figure 4G). These observed sex differences are not due to dose-dependent effects of βARKnt expression, as βARKnt-Flag expression was similarly varied across the male and female groups (Figure 4H). These data show a robust sex difference in VDAC1 and VDAC3 protein expression that may contribute to baseline sex differences in our tissue respiration measures. Further, they demonstrate that both control and βARKnt females exhibit alterations in these key mitochondrial protein expression levels with pressure overload stress.

Glucose tolerance tests (GTT) and insulin tolerance tests (ITT) 4 weeks after TAC in males (Figure 4I, S5I) and females (Figure 4J, S5J) or 14 weeks after TAC in females (Figure 4K, S5K) were used to assess systemic metabolic performance. Quantification of area under the curve from these tests showed that while male control and βARKnt, Sham and TAC mice exhibited no differences in glucose handling, it was enhanced in βARKnt TAC females after acute pressure overload (Figure 4L). Glucose tolerance was further altered in βARKnt Sham and more so TAC females after chronic pressure overload, wherein blood glucose levels never reached the level of elevation seen in control females and maintained a rapid return to baseline fasted levels (Figure 4M). While insulin sensitivity trended towards higher in females in general compared to males, this value was not significantly altered in βARKnt male or female, Sham or TAC mice compared to controls at 4 weeks after TAC (Figure 4N) and was only shifted in βARKnt TAC females 14 weeks after TAC with an impaired return towards fasted levels (Figure 4O). These data suggest enhanced glucose handling and insulin responses in βARKnt females compared to males that were further exaggerated during the response to chronic pressure overload stress.

### 2.5 βARKnt female mice exhibit significant differences in serum metabolite and metabolic pathway prevalence during acute and chronic pressure overload stress

Untargeted serum metabolomics was performed by Dr. Isaac Ampong in the Lerner Research Institute Proteomics & Metabolomics Core (Director, Dr. Belinda Willard) according to their standard practices and as detailed in the Materials and Methods section below. This analysis was performed on 48 serum samples obtained from NLC and βARKnt females with (TAC) or without surgery (Sham) at two time points (4 weeks and 14 weeks) to compare the metabolite changes between groups. Untargeted metabolomics data analysis identified a total of 7,821 metabolite features with 3508 MS1-level matches and 75 MS2-level matched to MSP spectral databases (Table 1). Data were analyzed using the multivariate statistical methods of principal component analysis (PCA) and orthogonal partial least squares discriminant analysis (OPLS-DA) to identify the metabolite differences between groups at given post-surgery time points. The OPLS-DA scores plots showed a strong separation of metabolome profiles in all comparisons (Figure 5A-D, top left). The top 15 most impacted metabolites from the multivariate analysis at 4 weeks after Sham (Figure 5A, top right), 4 weeks after TAC (Figure 5B, top right), 14 weeks after Sham (Figure 5C, top right), and 14 weeks after TAC (Figure 5D, top right) surgery are depicted. Volcano plots show the differential metabolites in the βARKnt versus NLC group for 4 weeks after Sham (Figure 5A, bottom), 4 weeks after TAC (Figure 5B, bottom), 14 weeks after Sham (Figure 5C, bottom), and 14 weeks after TAC (Figure 5D, bottom) surgery, respectively, demonstrating the strongest differences in metabolite abundance in βARKnt 14 weeks after TAC. Pathway enrichment mapping of the full metabolomics dataset 4 weeks after Sham (Figure S6A), 4 weeks after TAC (Figure S6B), 14 weeks after Sham (Figure S6C), and 14 weeks after TAC (Figure S6D) surgery with SMPDB using MetaboAnalyst revealed differential metabolic pathway alterations as a factor of genotype and time exposed to pressure overload stress. This analysis revealed overrepresentation of metabolites impacting 8 metabolic pathways for βARKnt vs NLC Sham at 4 weeks after surgery, 0 pathways for βARKnt vs NLC TAC at 4 weeks, 3 pathways for βARKnt vs NLC Sham at 14 weeks, and 14 metabolic pathways for βARKnt vs NLC TAC at 14 weeks after surgery (Table S1). Anywhere from 1-16 metabolite hits within a given metabolic pathway were overrepresented. These data were compared to a database of similar samples and analysis of that metabolite containing pathway, thus demonstrating a greater abundance of these 1-16 metabolites than could be expected by chance variation for each given sample type and suggesting a significant remodeling of the serum metabolome by cardiomyocyte βARKnt expression. The most affected pathways were at 14 weeks after TAC surgery and were related to amino acid metabolism, lipid metabolism, oxidative stress, and glucose degradation, suggesting the greatest impact to the serum metabolome during chronic pressure overload stress. Further interrogation and follow-up targeted metabolomics will hopefully provide key information toward elucidating the mechanism by which cardiac βARKnt is able to alter cardiac and systemic metabolism and function in these mice.

**Figure 5:**
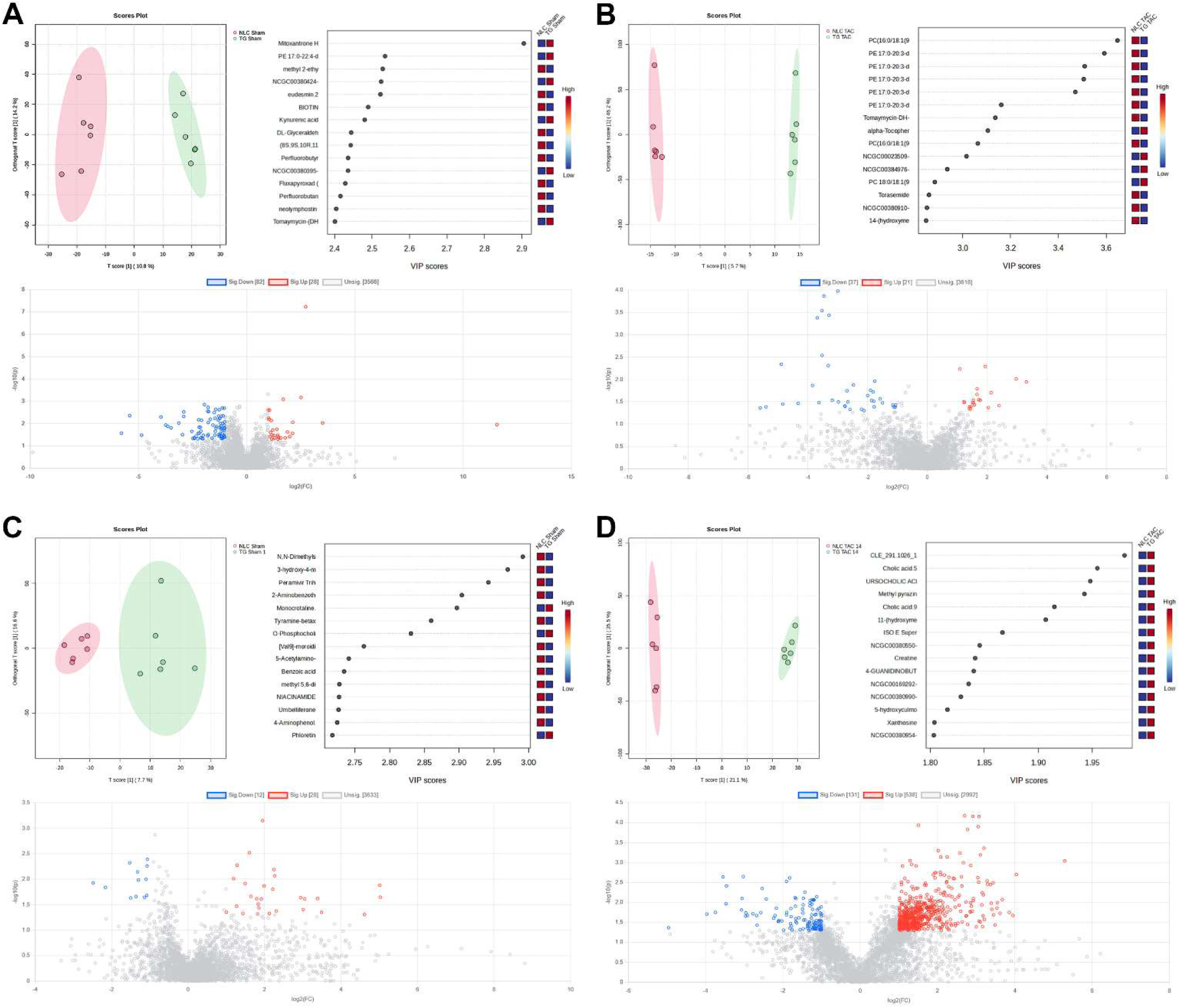
Untargeted serum metabolomics demonstrates strong differences in metabolite prevalence as a factor of genotype and time exposed to pressure overload stress. Untargeted metabolomics analysis of non-transgenic littermate control (NLC) and TgβARKnt (TG) serum **(A)** 4 weeks after Sham surgery, **(B)** 4 weeks after TAC surgery, **(C)** 14 weeks after Sham surgery, and **(D)** 14 weeks after TAC surgery showing (left top) orthogonal partial least squares discriminant analysis (OPLS-DA), (right top) the 15 most impacted metabolites, and (bottom) volcano plots from the multivariate analysis between NLC and βARKnt mice at these time points. The volcano plots show the individual statistically significant ions upregulated in the βARKnt mice compared to their respective controls depicted in red dots and downregulated in blue dots. The x- axis represents log2 fold-change which shows the direction of the change (negative scale is downregulated and positive scale is upregulated) in metabolite levels while the y-axis is the negative log of the p-value (FDR adjusted) showing the significance of the change.

**Table 1:** Untargeted Metabolomics Data Summary.

|  | POS ESI | NEG ESI | Total |
| --- | --- | --- | --- |
| **Number of Features** | 7287 | 534 | 7821 |
| **Number of MS1 Level Matches** | 3508 | 194 | 3702 |
| **Number of MS2 Level Features Matched to Reference Spectra** | 61 | 14 | 75 |

## 3. Discussion

In this study we investigated whether cardiomyocyte-specific transgenic expression of βARKnt in female mice would influence cardiac remodeling and the progression to heart failure after pressure-overload (TAC). Cardiac-specific TgβARKnt female mice exhibited enhanced fractional shortening compared to controls and an equivalent adaptive hypertrophy after acute TAC, though accompanied by significant left atrial hypertrophy. Chronic TAC exposure revealed a still elevated fractional shortening, though more robust in Sham than TAC animals, with a continuous hypertrophic thickening of the septal wall and now more robust increase in heart weight and left atrial weight compared to controls. Interestingly, this was accompanied by an increase in lung weight and a decrease in exercise tolerance indicating a transition to heart failure. While the increase in myocyte cross-sectional area was proportional with controls compared to their own Sham, fibrotic area was significantly elevated in both the ventricular myocardium and left atrium, and numerous small cells were evident amongst the cardiomyocytes, suggesting that interstitial fibrosis and infiltration or expansion of other cell types play a major role in the increase in wall thickness observed even during diastole. Analysis of 8 weeks old males and females showed that some, but not all, of these changes are present or beginning even at this early time point. Tissue respiration measures using ETC-specific substrates and inhibitors revealed sex- and genotype-driven alteration in mitochondrial function with the greatest inhibition occurring in βARKnt females 4 weeks after TAC. This was accompanied by similar changes in mitochondrial DNA and resident protein expression, suggesting potential alterations in mitochondrial content and function. Further, unlike control or βARKnt males with Sham or TAC surgery, both 4 and 14 weeks after surgery the βARKnt females showed alterations in systemic glucose responsiveness with a diminished increase in blood glucose yet effective return to baseline. While serum metabolomics offers some insights into the underlying systemic changes, future work is needed to reveal the underlying mechanisms.

Our results indicate differences in hypertrophy when βARKnt females were exposed to acute and chronic pressure overload stress. At baseline βARKnt female mice have elevated fractional shortening and hypertrophy, indicating an innate growth response to βARKnt expression. Both βARKnt and non-transgenic mice exhibit proportional left ventricular hypertrophic growth in response to acute TAC, but the βARKnt mice show differential left atrial remodeling, indicated by a significant increase in myocyte and fibrotic area, that is unique. These data suggest a normal physiological hypertrophic response to acute stress in the ventricles, despite the preexisting enlargement, which is a typical compensatory mechanism that helps maintain cardiac function in response to injury or stress. Yet, there is potential pathological remodeling in the βARKnt atria that could lead to the development of arrhythmias or heart failure. Additionally, baseline fibrosis and macrophage infiltration in βARKnt hearts suggests a preexisting vulnerability that may become exacerbated during chronic pressure overload stress. With chronic exposure to pressure overload, βARKnt female mice experienced adverse, asymmetrical ventricular remodeling with increased fibrosis and immune cell infiltration, suggesting a further transition to pathological remodeling, despite their preserved ejection fraction. In addition, our data showed pulmonary congestion and exercise intolerance in βARKnt mice after chronic TAC. Together these data suggest that βARKnt females develop a restrictive cardiomyopathy or HFpEF-like heart failure.[16, 17] This contrasts with control females that exhibited a more physiological hypertrophy and cardioprotection. This is also quite distinct from our male study, where control siblings transitioned from adaptive to maladaptive remodeling, decompensating into dilated cardiomyopathy or heart failure with reduced ejection fraction, while βARKnt males exhibited a more physiological hypertrophy and cardioprotection.

Pathological hypertrophy in heart failure is characterized by decreases in energy production and mitochondrial function alongside shifts in cardiac and systemic metabolism that contribute to disease progression.[18]. Impaired tissue respiration after TAC in βARKnt females, but not males, suggests compromised mitochondrial bioenergetics and inefficient ATP generation, which further supports the advancement of pathological hypertrophy. Interestingly, these Oroboros data in permeabilized tissue sections were distinct from previous Seahorse analysis of freshly dissociated contractile adult cardiomyocytes. βARKnt male myocytes fed cell level substrates showed enhanced spare respiratory capacity and ATP production compared to controls in the presence of both fatty acid substrate palmitate and glucose, showing increased efficiency in metabolite usage.[9] Female cardiomyocytes showed no difference. Together, these data suggest that βARKnt male myocytes may be more efficient in taking up substrates at the cell membrane to feed an otherwise intact ETC, whereas βARKnt females may not exhibit distinct differences in cellular metabolite uptake but have a decrease in mitochondrial content or health that impairs their function. This agrees with our previous proteomic work which detected a functional interaction of βARKnt with Akt substrate of 160 kDa (AS160), a key regulator of insulin-stimulated metabolite uptake.[9] Future proteomic analysis of βARKnt interacting partners in females will determine if they play a key role in the declined cardiomyocyte metabolism. A more in-depth analysis of cell membrane vs mitochondrial substrate uptake and mitochondrial content and structure will also provide a greater perspective on sex and genotype driven differences in mitochondrial function at baseline and following cardiac stress, and the role they play in cardiomyocyte metabolism and heart failure progression.

Systemic glucose tolerance is often impaired in heart failure. Particularly in its chronic stages, the heart will shift from fatty acid oxidation to glucose utilization, but systemic insulin resistance reduces glucose uptake in cardiac and skeletal muscle.[19, 20] This leads to hyperglycemia and further exacerbates heart failure symptoms, contributing to worse outcomes and disease progression. Although all males and control and βARKnt Sham female mice showed no differences in glucose handling, it was improved in βARKnt TAC females acutely after TAC and more so in βARKnt Sham and TAC female mice after chronic TAC. However, insulin sensitivity was not altered in male or female transgenic mice after acute pressure overload, representing a disconnect between systemic insulin sensitivity and cardiac metabolism.

Emerging research indicates that women experience distinct clinical manifestations and progression of heart failure compared to men, notably differences in heart failure with preserved ejection fraction (HFpEF), accounting for about two thirds of HFpEF cases, and associated comorbidities such as hypertension and obesity.[21, 22] Additionally, metabolic syndrome, linked to cardiovascular disease, varies by sex and yet female subjects are also underrepresented in cardiometabolic studies.[23, 24] Our previous studies highlight significant sex differences in cardiac remodeling and function in male and female βARKnt mice under metabolic stress, suggesting unique cardiac regulatory mechanisms between sexes.[25, 26] We investigated the effects of cardiomyocyte-specific expression of an amino-terminal peptide of GPCR kinase 2 (βARKnt) versus non-transgenic littermate controls (NLCs) on high-fat diet (HFD).[25, 26] Males showed enhanced cardiometabolic signaling, cardioprotection, metabolic flexibility, and improved white adipose tissue (WAT) health.[25] Females exhibited decreased cardiometabolic signaling, maladaptive remodeling, cardiac dysfunction, and diminished lipid accumulation and activation in brown adipose tissue (BAT), despite demonstrating enhanced systemic metabolism.[26] These data highlighted sex and stress specific systemic metabolic responses that led to different pathways of heart failure and revealed a novel and sex-dependent cardiac-mediated regulation of white (male) and brown (female) adipose tissue lipolysis and function during metabolic stress. Herein, control females exhibited a more physiological and cardioprotected response to pressure overload, while βARKnt females exhibited robust restrictive cardiomyopathy and heart failure with symptoms more akin to heart failure with preserved ejection fraction (HFpEF). Further, in response to metabolic stress, male and female βARKnt mice exhibit separate systemic metabolic profiles compared to their controls, between sexes, and as a factor of time exposed to HFD, with unique up- and down-regulated compounds. Herein, we also performed serum metabolomics analyses that further suggests potential associations between βARKnt expression and serum metabolite abundances that may underlie the pathophysiological responses. Comparison of these data may provide new avenues for understanding the underlying sex-linked mechanisms of cardiac remodeling and metabolic dysregulation. Future targeted metabolomics and metabolic tracer analyses will uncover biomarkers and pathways involved in the observed phenotype and evaluate the influences of βARKnt. Overall, these data emphasize the complex relationship between cardiac remodeling and metabolic adaptations in βARKnt transgenic mice, pointing to critical sex differences that warrant further studies on understanding the divergent pathophysiology and signaling in females during heart failure.

In summary, these results highlight how the expression of the short N-terminal domain of GRK2, βARKnt, plays a significant role in the sex-specific progression of cardiovascular disease and heart failure. Previous studies have suggested that female hearts often exhibit more robust adaptive mechanisms in response to acute stress, as evidenced by preserved ejection fractions and hypertrophic responses, as seen in our control females.[27, 28] The transition to heart failure observed in the βARKnt mice, despite maintained fractional shortening, echoes reports that suggest women may experience different pathways toward heart failure, including adverse remodeling and inflammation.[16, 29] The increased fibrosis and immune cell infiltration noted in βARKnt females are also consistent with literature identifying inflammation as a key player in heart failure progression.[30, 31] Some studies have shown that estrogen protects against diastolic dysfunction by enhancing mitochondrial function and insulin sensitivity and reducing oxidative stress.[28, 32, 33]. However, our findings differ in that the female βARKnt mice exhibit impaired mitochondrial function alongside enhanced insulin sensitivity, suggesting that the predominant phenotype, especially the unique left atrial enlargement, arises from a distinct cardiometabolic regulatory role of βARKnt. Future studies will examine hormonal influences and other sex-dependent factors that may be responsible for the observed cardioprotection in control females versus asymmetric diastolic left ventricular hypertrophy and heart failure in βARKnt females. Collectively, these results underscore the sex-specific aspects of cardiovascular disease and stress the need to investigate the underlying pathophysiological mechanisms. Ongoing βARKnt work uses this peptide as a tool to investigate the sex differences in protein interactions governing normal, adaptive, and maladaptive cardiovascular signaling to identify new therapeutic targets for the treatment of cardiovascular and cardiometabolic disease that may influence therapeutic strategies and outcomes in heart failure management.

## 4. Materials and Methods

### 4.1 Study Design & Experimental Animals

This study will focus on the use of 8–10-week-old female laboratory mice for all proposed experiments. Since results in females were quite divergent and due to the nature of the divergence, the focus of this manuscript is distinct from the previously published male study.[9] The lines used in this proposal are all driven by the cardiac-restricted promoter α-myosin heavy chain (αMHC) and are on a C57BL/6J background. Cardiac-specific transgenic (Tg) mice expressing a flag-tagged βARKnt peptide were generated as described previously [7]. Briefly, the cDNA that encodes bovine GRK2 residues 50- 145 was cloned into a vector driven by the αMHC promoter with a carboxyl-terminal flag-tag. All transgenic mouse studies include equal numbers of non-transgenic littermate control (NLC) mice for all planned experiments. Statistical powering was performed using the G*Power 3.1.9.2 software from the University of Dusseldorf for power analysis and estimation of sample size. Based on these calculations our target was a minimum of 9 animals (powered for indices of trans-aortic constriction) per group to attain statistical significance, with a preference for 10-15.

Cohorts of paired transgenic and non-transgenic littermate controls were randomly assigned an analytical group at each takedown (following 6 hours 4°C cold challenge), generally completing the n value needed for histology and then biochemistry. Regarding heart samples, whole hearts were either collected for 4 chamber histology as described below, or for biochemistry. For biochemistry, the same portion of the LV septum was removed from each for RNA analysis and the rest of the heart was used for Western blotting, such that the same exact samples were used for Western blotting and RT-PCR. All animals and resulting samples were monitored by mouse number only until data quantification was complete and then decoded by gene expression and surgical group for statistical analysis, meaning all data analysis was blinded. All results were substantiated by repetition. Data were only excluded if their validity was undermined by the condition of the animal or cells prior to or during the experiment, such as loss of the specimen. All animal procedures were carried out according to National Institutes of Health Guidelines on the Use of Laboratory Animals and approved by the Animal Care and Use Committees of Temple University and the Lerner Research Institute (LRI).

### 4.2 In vivo TAC Model & Echocardiography

Transverse aortic constriction (TAC) was performed as described previously [3, 34]. Echocardiography was performed on NLC and βARKnt mice using the Vevo 2100 imaging system from VisualSonics as described [35]. Briefly, two-dimensional echocardiographic views of the mid-ventricular short axis were obtained at the level of the papillary muscle tips below the mitral valve. M-mode measurements were determined at the plane bisecting the papillary muscles according to the American Society of Echocardiography leading edge method. To measure global cardiac function, echocardiography was performed at 8-10 weeks of age prior to TAC and at 2 and 4, or 2, 4, 6, 8, 10, 12 and 14 weeks post-TAC. Pressure gradients were determined 1 week later by pulsed-wave Doppler echocardiography of the transverse aorta and mice with gradients over 50 mmHg were used. Subgroups of hearts were harvested at 4 and 14 weeks after TAC for analyses of structure, fibrosis, and biochemistry.

### 4.3 Exercise Tolerance Testing

Pairs of mice were placed onto the specialized tracks of an LE8709 small animal forced exercise treadmill (Panlab) with the speed set to 5 cm/s. Mice were run at 5 cm/s for 5 minutes, then at the 5^th^ minute, the speed was increased to 10 cm/s, and at the 6^th^ minute and every minute after, the speed was increased by another 5 cm/s. The machine display shows 5 parameters that include distance, shock time, shock number, speed, and time. Shock time was monitored throughout the running duration to ensure that the mice were maintaining the ability to run. Each mouse was removed from the treadmill track when shock time approached 15 seconds and distance, speed, and time spent running were recorded for each mouse at this endpoint. If both mice were approaching 15s simultaneously, then the experiment was stopped and record the distance, speed, and time parameters were recorded.

### 4.4 Histological Sectioning and Staining

Subgroups of hearts were harvested at 4 and 14 weeks post-TAC for analyses of structure, fibrosis, and biochemistry. Trichrome staining was performed as previously described [36]. Briefly, mice were euthanized and following cardioperfusion, hearts were fixed for 1-3 days in 4% paraformaldehyde at 4°C. Hearts were dehydrated and paraffinized using a Microm STP 120 from ThermoFisher Scientific, embedded in paraffin using a HistoStar apparatus (ThermoFisher), and sectioned (4-6 micron) using a Microm HM 325 (ThermoFisher). Trichrome staining was performed as previously described.[37] Heart sections were de-paraffinized and stained with Weigert’s iron hematoxylin and Masson Trichrome (Sigma-Aldrich) according to the manufacturer’s instructions. Sections were then scanned on an Aperio AT2 digital whole slide scanner (Leica Biosystems) and quantified using the Positive Pixel Count v9 algorithm in the Aperio ImageScope analysis software (Leica).

Alternatively, tissue sections were de-paraffinized and re-hydrated according to the Trichrome staining protocol, but following the wash with de-ionized water (dH20) heart sections were washed 3 times 5 minutes with 1x PBS followed by staining with Alexa Fluor 488-conjugated wheat germ agglutinin (WGA, 1:10 in 1xPBS) (Invitrogen) for 1 hour at room temperature in a humidified chamber in the dark. Sections were again washed 3 times 5 minutes with 1x PBS followed by mounting with coverslips using Fluoromount-G mounting media containing DAPI nuclear stain (Southern Biotech). CD68 staining was performed according to the Cell Signaling protocol for antibody #97778. Briefly, tissue sections were de-paraffinized and re-hydrated according to the Trichrome staining protocol, but following the water wash, sections underwent antigen retrieval in 10 mM sodium citrate buffer, pH 6.0, at sub-boiling temperatures (95-98°C) for 10 minutes. After cooling to room temperature, tissue sections were washed 3 times 5 minutes with dH2O, incubated with 3% hydrogen peroxide for 10 minutes, washed 2 times 5 minutes with dH2O followed by 1 times 5 minutes in Tris Buffered Saline with Tween 20 (TBST) wash buffer. Next, tissue sections were blocked with 5% Normal Goat Serum (Cell Signaling Technology, CST) in TBST and incubated with rabbit monoclonal CD68 (E3O7V) diluted in SignalStain Antibody Diluent (CST) at 4°C overnight. Sections were then washed 3 times 5 minutes in wash buffer, incubated with SignalStain Boost Detection Reagent (CST) for 30 minutes in a humidified chamber, washed 3 times 5 minutes with wash buffer and incubated with SignalStain DAB reagent (CST) until acceptable staining intensity (∼10 minutes), dehydrated with increasing ethanol concentrations followed by xylene, and mounted with Permount. Sections were then scanned on an Aperio AT2 digital whole slide scanner (Leica Biosystems) and quantified using the Positive Pixel Count v9 algorithm in the Aperio ImageScope analysis software (Leica).

### 4.5 RNA Isolation and Semi-quantitative PCR

RNA isolation and analysis was performed as previously described [3]. Total RNA isolation was performed using TRIzol reagent (Life Technologies) and a VCX 130PB ultrasonic processor from Sonics & Materials Inc (Newtown, CT) for homogenization. After RNA isolation, cDNA was synthesized from 1mg of total RNA using the iScript cDNA Synthesis Kit from Bio-Rad Laboratories (Hercules, CA). Semi-quantitative PCR was conducted on cDNA using SYBR Green (Bio-Rad) and 150nM of gene-specific oligonucleotides for*18S*, *ANP*, *βMHC, Bcl-xL, CHOP, TGFβ, and TNFα* on a CFX96 Real Time System with BioRad CFX Manager 2.1 software (Bio-Rad). Quantitation was established by comparing 18s mRNA, which was similar between groups, for normalization and compared using the ΔΔCt method.

### 4.6 Oroboros Oxygraph Assessment of Permeabilized Cardiac Myofiber Respiration

Permeabilized LV fibers were used to determine mitochondrial respiratory capacity, using a polarographic oxygen sensor (Oxygraph-2k, Oroboros, Innsbruck, Austria). Studies were performed in carefully weighed fresh LV muscle from 5-6 mice in each group. Heart muscle fibers (∼3mg) were dissected and separated with fine tipped forceps. Chemical permeabilization was performed by incubation with saponin (50 μg/ml) in 2ml ice cold biopsy buffer solution BIOPS (CaK2EGTA 2.77 mM, K2EGTA 7.23 mM, Na2ATP 5.77 mM, and sodium phosphocreatine 15 mM, MgCl2.6H2O 6.56 mM, taurine 20 mM, imidazole 20 mM, dithiothreitol 0.5 mM, 2-N morpholino ethane sulfonic acid hydrate (MES) 50 mM, free calcium 0.1 μM, free magnesium 1 mM, and Mg ATP 5 mM) at pH 7.1[13] for 20 min at 4°C and washed for10 min in BIOPS, followed by washing in miRO5 buffer (EGTA 0.5 mM, MgCl2 6H2O 3 mM, K-lactobionate 60 mM, taurine 20 mM, K phosphate 10 mM, HEPES 20 mM, sucrose 110 mM, bovine serum albumin (essentially FA free, fraction V) 1g/l). Samples (1.5-2 mg wet weight) were transferred into a chamber of the Oroboros Oxygraph containing air-saturated assay medium MiRO6 (MiRO5 plus 280 units/ml catalase) at 37°C. Oxygen concentration and flow rates were recorded at 2- second intervals, after calibration of the oxygen sensors and instrument background corrections. Standard substrate, uncoupler, and inhibitor titration (SUIT) protocols were used to quantify mitochondrial function.[13–15] Oxygen flux in heart muscle fibers was assessed after sequential additions of substrates, ADP (2.5 mM), uncoupler, and inhibitors. Mitochondrial substrates pyruvate (5mM), malate (2 mM), followed by ADP (2.5 mM) and glutamate (10mM) were added to measure complex I function. Succinate (50mM) was then added as a complex II substrate. Maximal oxidative capacity, or maximum respiration, was measured by the addition of uncoupler-FCCP (1ul of 1mM FCCP), followed by rotenone (0.5 uM) to inhibit electron flow across complex I. Antimycin A (2.5 uM) complex III inhibitor was then added to determine the non-mitochondrial residual oxygen consumption rate. The uncoupled complex IV oxidation rate was calculated using an artificial electron donor to complex IV N,N,N′,N′-tetramethyl-p-phenylenediamine (0.5mM TMPD) and ascorbate (2mM) followed by sodium azide (100 m M). When needed, reoxygenation was performed by titration of a solution of 200mM hydrogen peroxide into the chamber.

All data were expressed as oxygen consumption rates in pmol/s normalized to the wet weight of tissue to allow for comparisons across experiments. Data were analyzed using DatLab2 from Oroboros Instruments. Oxidation coupled to phosphorylation was quantified in response to substrates for complexes I and II and ADP. Mitochondrial respiration was corrected for oxygen flux due to instrumental background, and for residual oxygen consumption (ROX) after inhibition of Complexes I and III, with rotenone and antimycin A, respectively. ROX was subtracted from oxygen flux as a baseline for all respiratory states.

### 4.7 Glucose and Insulin Tolerance Tests (GTT, ITT)

GTTs and ITTs were done as described.[38] In brief, GTT was used as a readout for glucose stimulated insulin secretion, and as an indication of systemic glucose metabolism. 4- or 14-weeks after Sham or TAC surgery, NLC and βARKnt mice were fasted for 6 hours and then injected intraperitoneally with a sterile dextrose solution (2g/kg body weight). A drop of blood was collected by nicking the tail, and blood glucose levels were measured using a glucometer (Serial No. 10S00007552, Clarity Diagnostics, Boca Raton FL) at 0, 15, 30, 60, 90 and 120 minutes. Similarly, for ITT, 4- and 14-weeks after surgery, NLC and βARKnt mice were fasted for 4 hours and injected intraperitoneally with human recombinant insulin (0.75U/kg body weight). A drop of blood was collected by nicking the tail, and blood glucose levels were measured at 0, 15, 30, 60, 90 and 120 minutes as a standard measurements of systemic glucose uptake.

### 4.8 Untargeted Metabolomics

Forty-eight serum samples were prepared for untargeted metabolomics by diluting 50µl of each sample with 400µl of chilled extraction solvent (Acetonitrile: methanol: formic acid (75, 25, 0.2, v/v/v)) containing 4 internal standards as described.[26] The samples were then centrifuged at 14,000g for 20 minutes to precipitate out the protein pellet. The supernatants were collected into new tubes and dried by speedvac overnight. Dried tubes containing samples were reconstituted with 100 µl of reconstitution solution (95:5, water: acetonitrile). Samples were vortexed, centrifuged and subjected to Liquid Chromatography-Mass Spectrometry (LC-MS) analysis. 2 µL from each sample was taken and pooled and this pooled QC sample was analyzed in the beginning and ending, and at every 10th injection. In addition, MS2 level data were collected on representative control and treated samples in this study.

#### Liquid Chromatography-Mass Spectrometry (LC-MS)

Untargeted metabolomics using reverse phase chromatography were performed by injecting 6 µL of each sample on a Thermo Accucore Vanquish C18 column with dimensions 100 x 2.1 mm, 1.5 µm particle size (Thermo P/N 27101-102130) at 60°C coupled to a Thermo Vanquish UHPLC by gradient elution where mobile phase **A** is 0.1% formic acid in water and mobile phase **B** is 0.1% formic acid in acetonitrile.[26]

The Orbitrap Q Exactive HF was operated in positive and negative electrospray ionization modes in different LC-MS runs over a mass range of 56-850 Da using full MS at 120,000 resolution. Data dependent acquisitions (DDA) on the pooled representative QC samples include MS full scans at a resolution of 120,000 and HCD MS/MS scans taken on the top 5 most abundant ions at a resolution of 30,000 with dynamic exclusion of 40.0 seconds and the apex trigger set at 2.0 to 4.0 seconds. The resolution of the MS2 scans were taken at a stepped NCE energy of 10.0, 20.0 and 30.0.

#### Data Processing and Annotation

Data was processed using MSDIAL [39] (v.4.92) for feature detection, identification and alignment using parameters optimized for data acquired on an Orbitrap mass spectrometer. MS1 and MS2 were set to profile mode in both positive and negative ionization modes. Peak detection of MS1 and MS2 spectra were set to tolerances of 0.01 Da and 0.025 Da, respectively, over a mass range of 56-850 m/z with minimum peak width and height of 5 and 1,000,000. Annotation was done using publically available libraries from MassBank of North America (MoNA) containing 13,303 unique compounds (positive mode) [40] and MSDIAL Metabolomics MSP Spectral Kit [41] containing 12,879 unique compounds (negative mode) with 80% identification cut-off score.

#### Statistical Analysis

Spectral features from MSDIAL processed data were further analyzed via MetaboAnalyst 5.0 (www.metaboanalyst.ca) [42]. Zero/missing values were replaced by 1/5 of the minimum peak height over all samples, normalized by sum normalization, log transformed and autoscaled prior to downstream statistical analysis. A raw p-value less than 0.05 was chosen for significance. All p-values were corrected via the Benjamini-Hochberg procedure for false discovery rate (FDR) with a threshold of 0.05. Fold change analysis was performed. Multivariate principal component analysis (PCA) and hierarchical clustering were performed for understanding metabolite variation and expression patterns between groups. Statistical analysis was performed by comparing either the TG Sham to NLC Sham group or TG TAC to NLC TAC group for each time point. In addition, time series analysis was performed using the whole dataset for Sham or TAC with genotype (NLC and TG) controlled as a covariate.

#### Pathway Mapping

Pathway enrichment analysis was performed to identify top altered pathways for putatively identified metabolites in the dataset. In this analysis the Pathway-associated metabolite sets (SMPDB) database, which consists of 99 pathways from the Small Molecule Pathway Database[43] was used to map the putatively identified ions to various pathways.

### 4.9 Statistical Analysis

All values in the text and figures are presented as mean ± standard error of the mean (SEM) of independent experiments for given size N. Statistical significance was determined by one-way analysis of variance (ANOVA) with repeated measures (where appropriate and as indicated) and Tukey post-hoc test for multiple pairwise comparisons, or two-way ANOVA with repeated measures and Bonferroni’s post-hoc test for comparisons at a given time point. For all statistical tests, a p value of <0.05 was considered significant. Analyses were performed using Graph Pad Prism 9 software (Graph Pad).

## Supporting information

NT Female TAC Paper Supplement

## Author Contributions

KMB and SMS designed the experiments. KMB, IHM and JG managed animal husbandry. KMB led experiments performed and analyzed by KMB, IHM, ADI, MB, and KMV, with support from IHM and ADI. JHR and DRVW provided access to and data analysis and interpretation support for the Oroboros oxygraphy studies. IA performed the metabolomics isolation and analysis and provided an initial data report of key specs and outcomes. KMB, ADI and SMS prepared the manuscript.

## Acknowledgements

This work was supported by the National Institutes of Health R00 HL132882 to SMS, R01 HL160755 (SMS), and 1S10OD021561 to SNP.

## Abbreviations

GRK2: G protein-coupled receptor kinase 2
RGS: Regulator of G protein Signaling homology domain
βARKct: cardiac restricted expression of carboxyl-terminus domain of GRK2
βARKnt: cardiac restricted expression of amino-terminus domain of GRK2
βAR: beta-adrenergic receptor
TAC: transverse aortic constriction
NLC: non-transgenic littermate control
Tg: transgenic
LV: left ventricular
OXPHOS: mitochondrial oxidative phosphorylation
ETC: electron transport chain
SUIT: substrate uncoupler inhibitor titration
(LC-MS/MS): liquid chromatography-electrospray ionization-tandem mass spectrometry
HFrEF: heart failure with reduced ejection fraction
HFpEF: heart failure with preserved ejection fraction

## Reference List

1. Penela P, Murga C, Ribas C, Lafarga V, Mayor F, Jr. The complex G protein-coupled receptor kinase 2 (GRK2) interactome unveils new physiopathological targets. Br J Pharmacol. 2010;160(4):821–32. doi: 10.1111/j.1476-5381.2010.00727.x. PubMed PMID: 20590581; PubMed Central PMCID: PMCPMC2935989.

2. Sato PY, Chuprun JK, Ibetti J, Cannavo A, Drosatos K, Elrod JW, et al. GRK2 compromises cardiomyocyte mitochondrial function by diminishing fatty acid-mediated oxygen consumption and increasing superoxide levels. J Mol Cell Cardiol. 2015;89(Pt B):360–4. doi: 10.1016/j.yjmcc.2015.10.002. PubMed PMID: 26506135; PubMed Central PMCID: PMCPMC4689631.

3. Schumacher SM, Gao E, Cohen M, Lieu M, Chuprun JK, Koch WJ. A peptide of the RGS domain of GRK2 binds and inhibits Galpha(q) to suppress pathological cardiac hypertrophy and dysfunction. Sci Signal. 2016;9(420):ra30. Epub 2016/03/27. doi: 10.1126/scisignal.aae0549. PubMed PMID: 27016525; PubMed Central PMCID: PMCPMC5015886.

4. Koch WJ, Inglese J, Stone WC, Lefkowitz RJ. The binding site for the beta gamma subunits of heterotrimeric G proteins on the beta-adrenergic receptor kinase. J Biol Chem. 1993;268(11):8256–60. PubMed PMID: 8463335.

5. Koch WJ, Rockman HA, Samama P, Hamilton RA, Bond RA, Milano CA, et al. Cardiac function in mice overexpressing the beta-adrenergic receptor kinase or a beta ARK inhibitor. Science. 1995;268(5215):1350–3. doi: 10.1126/science.7761854. PubMed PMID: 7761854.

6. Chen M, Sato PY, Chuprun JK, Peroutka RJ, Otis NJ, Ibetti J, et al. Prodeath signaling of G protein-coupled receptor kinase 2 in cardiac myocytes after ischemic stress occurs via extracellular signal-regulated kinase-dependent heat shock protein 90-mediated mitochondrial targeting. Circ Res. 2013;112(8):1121–34. doi: 10.1161/CIRCRESAHA.112.300754. PubMed PMID: 23467820; PubMed Central PMCID: PMCPMC3908784.

7. Keys JR, Greene EA, Cooper CJ, Naga Prasad SV, Rockman HA, Koch WJ. Cardiac hypertrophy and altered beta-adrenergic signaling in transgenic mice that express the amino terminus of beta-ARK1. Am J Physiol Heart Circ Physiol. 2003;285(5):H2201–11. doi: 10.1152/ajpheart.00112.2003. PubMed PMID: 12869383.

8. Sterne-Marr R, Tesmer JJ, Day PW, Stracquatanio RP, Cilente JA, O’Connor KE, et al. G protein-coupled receptor Kinase 2/G alpha q/11 interaction. A novel surface on a regulator of G protein signaling homology domain for binding G alpha subunits. J Biol Chem. 2003;278(8):6050–8. doi: 10.1074/jbc.M208787200. PubMed PMID: 12427730.

9. Bledzka KM, Manaserh IH, Grondolsky J, Pfleger J, Roy R, Gao E, et al. A peptide of the amino-terminus of GRK2 induces hypertrophy and yet elicits cardioprotection after pressure overload. J Mol Cell Cardiol. 2021;154:137–53. Epub 2021/02/07. doi: 10.1016/j.yjmcc.2021.01.004. PubMed PMID: 33548241; PubMed Central PMCID: PMCPMC8101069.

10. Reza N, Gruen J, Bozkurt B. Representation of women in heart failure clinical trials: Barriers to enrollment and strategies to close the gap. Am Heart J Plus. 2022;13. Epub 2022/03/05. doi: 10.1016/j.ahjo.2022.100093. PubMed PMID: 35243454; PubMed Central PMCID: PMCPMC8890694.

11. Lam CSP, Arnott C, Beale AL, Chandramouli C, Hilfiker-Kleiner D, Kaye DM, et al. Sex differences in heart failure. Eur Heart J. 2019;40(47):3859–68c. Epub 2019/12/05. doi: 10.1093/eurheartj/ehz835. PubMed PMID: 31800034.

12. Siraki AG. The many roles of myeloperoxidase: From inflammation and immunity to biomarkers, drug metabolism and drug discovery. Redox Biol. 2021;46:102109. Epub 2021/08/30. doi: 10.1016/j.redox.2021.102109. PubMed PMID: 34455146; PubMed Central PMCID: PMCPMC8403760.

13. Pesta D, Gnaiger E. High-resolution respirometry: OXPHOS protocols for human cells and permeabilized fibers from small biopsies of human muscle. Methods Mol Biol. 2012;810:25–58. doi: 10.1007/978-1-61779-382-0_3. PubMed PMID: 22057559.

14. Davuluri G, Allawy A, Thapaliya S, Rennison JH, Singh D, Kumar A, et al. Hyperammonaemia-induced skeletal muscle mitochondrial dysfunction results in cataplerosis and oxidative stress. J Physiol. 2016;594(24):7341–60. doi: 10.1113/JP272796. PubMed PMID: 27558544; PubMed Central PMCID: PMCPMC5157075.

15. Kumar A, Davuluri G, Welch N, Kim A, Gangadhariah M, Allawy A, et al. Oxidative stress mediates ethanol-induced skeletal muscle mitochondrial dysfunction and dysregulated protein synthesis and autophagy. Free Radic Biol Med. 2019;145:284–99. Epub 2019/10/02. doi: 10.1016/j.freeradbiomed.2019.09.031. PubMed PMID: 31574345; PubMed Central PMCID: PMCPMC6910229.

16. Redfield MM. Heart Failure with Preserved Ejection Fraction. N Engl J Med. 2016;375(19):1868–77. doi: 10.1056/NEJMcp1511175. PubMed PMID: 27959663.

17. Mohammed SF, Hussain S, Mirzoyev SA, Edwards WD, Maleszewski JJ, Redfield MM. Coronary microvascular rarefaction and myocardial fibrosis in heart failure with preserved ejection fraction. Circulation. 2015;131(6):550–9. doi: 10.1161/CIRCULATIONAHA.114.009625. PubMed PMID: 25552356; PubMed Central PMCID: PMCPMC4324362.

18. Yang D, Liu HQ, Liu FY, Guo Z, An P, Wang MY, et al. Mitochondria in Pathological Cardiac Hypertrophy Research and Therapy. Front Cardiovasc Med. 2021;8:822969. Epub 2022/02/05. doi: 10.3389/fcvm.2021.822969. PubMed PMID: 35118147; PubMed Central PMCID: PMCPMC8804293.

19. Doehner W, Frenneaux M, Anker SD. Metabolic impairment in heart failure: the myocardial and systemic perspective. J Am Coll Cardiol. 2014;64(13):1388–400. Epub 2014/09/27. doi: 10.1016/j.jacc.2014.04.083. PubMed PMID: 25257642.

20. Muniyappa R, Montagnani M, Koh KK, Quon MJ. Cardiovascular actions of insulin. Endocr Rev. 2007;28(5):463–91. Epub 2007/05/26. doi: 10.1210/er.2007-0006. PubMed PMID: 17525361.

21. Humphries KH, Izadnegahdar M, Sedlak T, Saw J, Johnston N, Schenck-Gustafsson K, et al. Sex differences in cardiovascular disease - Impact on care and outcomes. Front Neuroendocrinol. 2017;46:46–70. Epub 2017/04/22. doi: 10.1016/j.yfrne.2017.04.001. PubMed PMID: 28428055; PubMed Central PMCID: PMCPMC5506856.

22. DeFilippis EM, Van Spall HGC. Is it Time for Sex-Specific Guidelines for Cardiovascular Disease? J Am Coll Cardiol. 2021;78(2):189–92. Epub 2021/07/10. doi: 10.1016/j.jacc.2021.05.012. PubMed PMID: 34238440.

23. Pradhan AD. Sex differences in the metabolic syndrome: implications for cardiovascular health in women. Clin Chem. 2014;60(1):44–52. Epub 2013/11/21. doi: 10.1373/clinchem.2013.202549. PubMed PMID: 24255079.

24. Mauvais-Jarvis F. Sex differences in metabolic homeostasis, diabetes, and obesity. Biol Sex Differ. 2015;6:14. Epub 2015/09/05. doi: 10.1186/s13293-015-0033-y. PubMed PMID: 26339468; PubMed Central PMCID: PMCPMC4559072.

25. Manaserh IH, Bledzka KM, Junker A, Grondolsky J, Schumacher SM. A Cardiac Amino-Terminal GRK2 Peptide Inhibits Maladaptive Adipocyte Hypertrophy and Insulin Resistance During Diet-Induced Obesity. JACC Basic Transl Sci. 2022;7(6):563–79. Epub 2022/07/13. doi: 10.1016/j.jacbts.2022.01.010. PubMed PMID: 35818501; PubMed Central PMCID: PMCPMC9270572.

26. Manaserh IH, Bledzka KM, Ampong I, Junker A, Grondolsky J, Schumacher SM. A cardiac amino-terminal GRK2 peptide inhibits insulin resistance yet enhances maladaptive cardiovascular and brown adipose tissue remodeling in females during diet-induced obesity. J Mol Cell Cardiol. 2023;183:81–97. Epub 2023/09/16. doi: 10.1016/j.yjmcc.2023.09.001. PubMed PMID: 37714510; PubMed Central PMCID: PMCPMC10591815.

27. Abubakar M, Saleem A, Hajjaj M, Faiz H, Pragya A, Jamil R, et al. Sex-specific differences in risk factors, comorbidities, diagnostic challenges, optimal management, and prognostic outcomes of heart failure with preserved ejection fraction: A comprehensive literature review. Heart Fail Rev. 2024;29(1):235–56. Epub 2023/11/24. doi: 10.1007/s10741-023-10369-4. PubMed PMID: 37996694.

28. Wei J, Duan X, Chen J, Zhang D, Xu J, Zhuang J, et al. Metabolic adaptations in pressure overload hypertrophic heart. Heart Fail Rev. 2024;29(1):95–111. Epub 2023/09/28. doi: 10.1007/s10741-023-10353-y. PubMed PMID: 37768435.

29. Beale AL, Meyer P, Marwick TH, Lam CSP, Kaye DM. Sex Differences in Cardiovascular Pathophysiology: Why Women Are Overrepresented in Heart Failure With Preserved Ejection Fraction. Circulation. 2018;138(2):198–205. Epub 2018/07/11. doi: 10.1161/CIRCULATIONAHA.118.034271. PubMed PMID: 29986961.

30. Adamo L, Rocha-Resende C, Prabhu SD, Mann DL. Reappraising the role of inflammation in heart failure. Nat Rev Cardiol. 2020;17(5):269–85. Epub 2020/01/24. doi: 10.1038/s41569-019-0315-x. PubMed PMID: 31969688.

31. Boulet J, Sridhar VS, Bouabdallaoui N, Tardif JC, White M. Inflammation in heart failure: pathophysiology and therapeutic strategies. Inflamm Res. 2024;73(5):709–23. Epub 2024/03/28. doi: 10.1007/s00011-023-01845-6. PubMed PMID: 38546848; PubMed Central PMCID: PMCPMC11058911.

32. Ventura-Clapier R, Piquereau J, Veksler V, Garnier A. Estrogens, Estrogen Receptors Effects on Cardiac and Skeletal Muscle Mitochondria. Front Endocrinol (Lausanne). 2019;10:557. Epub 2019/09/03. doi: 10.3389/fendo.2019.00557. PubMed PMID: 31474941; PubMed Central PMCID: PMCPMC6702264.

33. Iorga A, Cunningham CM, Moazeni S, Ruffenach G, Umar S, Eghbali M. The protective role of estrogen and estrogen receptors in cardiovascular disease and the controversial use of estrogen therapy. Biol Sex Differ. 2017;8(1):33. Epub 2017/10/27. doi: 10.1186/s13293-017-0152-8. PubMed PMID: 29065927; PubMed Central PMCID: PMCPMC5655818.

34. Martini JS, Raake P, Vinge LE, DeGeorge BR, Jr., Chuprun JK, Harris DM, et al. Uncovering G protein-coupled receptor kinase-5 as a histone deacetylase kinase in the nucleus of cardiomyocytes. Proceedings of the National Academy of Sciences of the United States of America. 2008;105(34):12457–62. doi: 10.1073/pnas.0803153105. PubMed PMID: 18711143; PubMed Central PMCID: PMC2527933.

35. Brinks H, Boucher M, Gao E, Chuprun JK, Pesant S, Raake PW, et al. Level of G protein-coupled receptor kinase-2 determines myocardial ischemia/reperfusion injury via pro- and anti-apoptotic mechanisms. Circ Res. 2010;107(9):1140–9. doi: 10.1161/CIRCRESAHA.110.221010. PubMed PMID: 20814022; PubMed Central PMCID: PMCPMC2966514.

36. Gao E, Lei YH, Shang X, Huang ZM, Zuo L, Boucher M, et al. A novel and efficient model of coronary artery ligation and myocardial infarction in the mouse. Circulation research. 2010;107(12):1445–53. doi: 10.1161/CIRCRESAHA.110.223925. PubMed PMID: 20966393; PubMed Central PMCID: PMC3005817.

37. Bledzka KM, Manaserh IH, Grondolsky J, Pfleger J, Roy R, Gao E, et al. A peptide of the amino-terminus of GRK2 induces hypertrophy and yet elicits cardioprotection after pressure overload. J Mol Cell Cardiol. 2021. Epub 2021/02/07. doi: 10.1016/j.yjmcc.2021.01.004. PubMed PMID: 33548241.

38. Manaserh IH, Maly E, Jahromi M, Chikkamenahalli L, Park J, Hill J. Insulin sensing by astrocytes is critical for normal thermogenesis and body temperature regulation. J Endocrinol. 2020;247(1):39–52. Epub 2020/07/23. doi: 10.1530/JOE-20-0052. PubMed PMID: 32698146; PubMed Central PMCID: PMCPMC7456332.

39. Tsugawa H, Cajka T, Kind T, Ma Y, Higgins B, Ikeda K, et al. MS-DIAL: data-independent MS/MS deconvolution for comprehensive metabolome analysis. Nat Methods. 2015;12(6):523–6. Epub 2015/05/06. doi: 10.1038/nmeth.3393. PubMed PMID: 25938372; PubMed Central PMCID: PMCPMC4449330.

40. MassBank of North America (MoNA) 2023. Available from: https://mona.fiehnlab.ucdavis.edu/

41. MS-DIAL metabolomics MSP spectral kit 2023. Available from: http://prime.psc.riken.jp/compms/msdial/main.html#MSP

42. Tsugawa H, Ikeda K, Takahashi M, Satoh A, Mori Y, Uchino H, et al. A lipidome atlas in MS-DIAL 4. Nat Biotechnol. 2020;38(10):1159–63. Epub 2020/06/17. doi: 10.1038/s41587-020-0531-2. PubMed PMID: 32541957.

43. Xia J, Psychogios N, Young N, Wishart DS. MetaboAnalyst: a web server for metabolomic data analysis and interpretation. Nucleic Acids Res. 2009;37(Web Server issue):W652–60. Epub 2009/05/12. doi: 10.1093/nar/gkp356. PubMed PMID: 19429898; PubMed Central PMCID: PMCPMC2703878.

