## Supplementary material for "Female Specific Restrictive Cardiomyopathy and Metabolic Dysregulation in transgenic mice expressing a Peptide of the Amino-Terminus of GRK2": NT Female TAC Paper Supplement

### Supplementary Appendix

#### Supplemental Figures

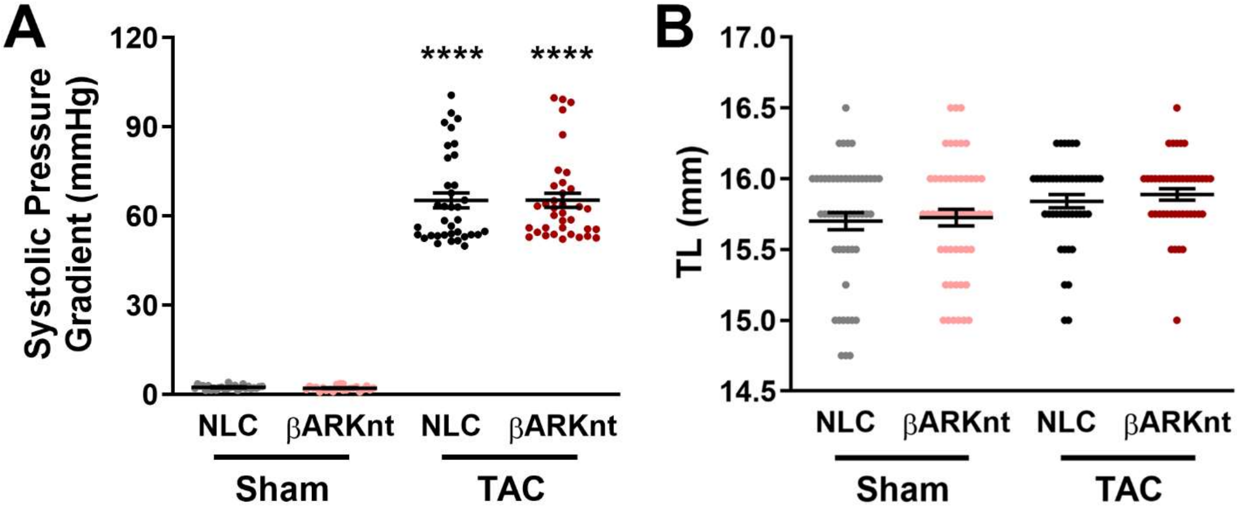


**Figure S1:** *Cardiomyocyte-specific βARKnt expression enhances cardiac fractional shortening and elicits baseline hypertrophy with adaptive pressure overload-induced hypertrophy.* (A) Measures of the systolic pressure gradient in non-transgenic littermate control (NLC) and TgβARKnt Sham and post-TAC mice 1 week after surgery. (B) Measures of tibia length in these animals 4 weeks after surgery. ****, $p < 0.0001$ by one-way ANOVA with Tukey post-hoc test relative to NLC Sham. n = 26-58 mice per group.

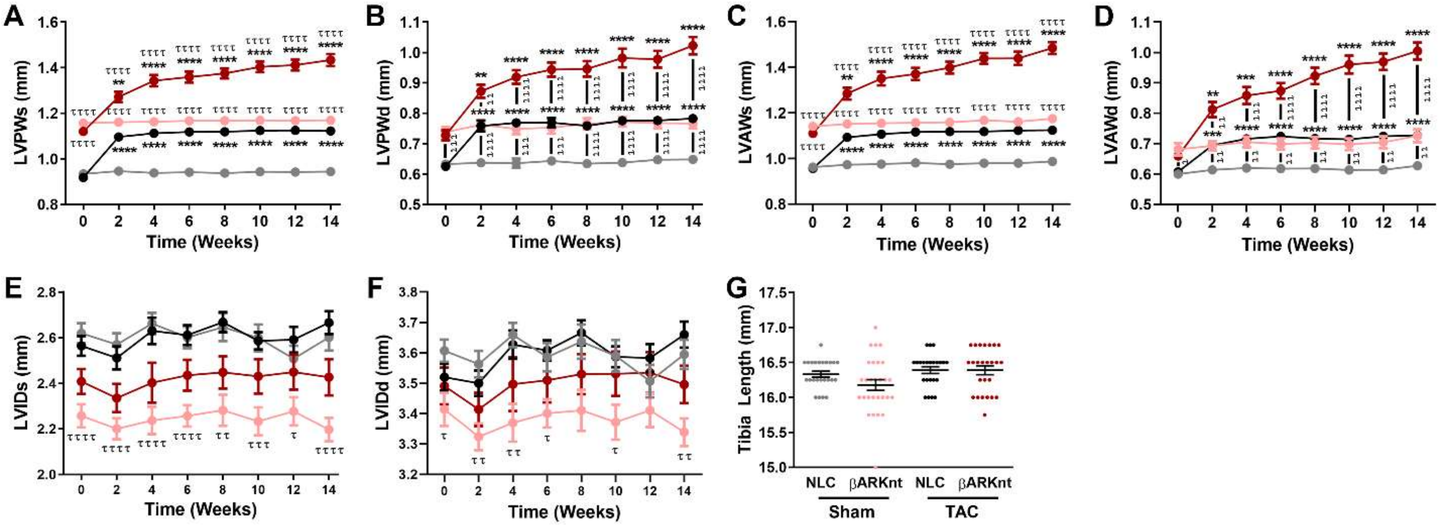


**Figure S2:** *Despite preserved cardiac function, βARKnt hearts exhibit asymmetric hypertrophic cardiomyopathy and heart failure after chronic pressure overload.* Serial measures of (A) left ventricular (LV) posterior wall thickness during systole (LVPWs), (B) LV posterior wall thickness during diastole (LVPWd), (C) LV anterior wall thickness during systole (LVAWs), (D) LV anterior wall thickness during diastole (LVAWd), (E) LV interior diameter during systole (LVIDs), and (F) LV interior diameter during diastole (LVIDd) in non-transgenic littermate control (NLC) and TgβARKnt mice 14 weeks after Sham or TAC surgery. **, $p < 0.01$; ***, $p < 0.01$; ****, $p < 0.0001$ by one-way ANOVA with Tukey post-hoc test relative to corresponding Sham. $^{\tau}$, $p < 0.05$; $^{\tau\tau}$, $p < 0.01$; $^{\tau\tau\tau}$, $p < 0.001$; $^{\tau\tau\tau\tau}$, $p < 0.0001$ by two-way ANOVA with repeated measures and Tukey post-hoc test relative to corresponding NLC. n = 16-20 mice per group. (B) Measures of tibia length in these animals 14 weeks after surgery. n = 16-29 mice per group.

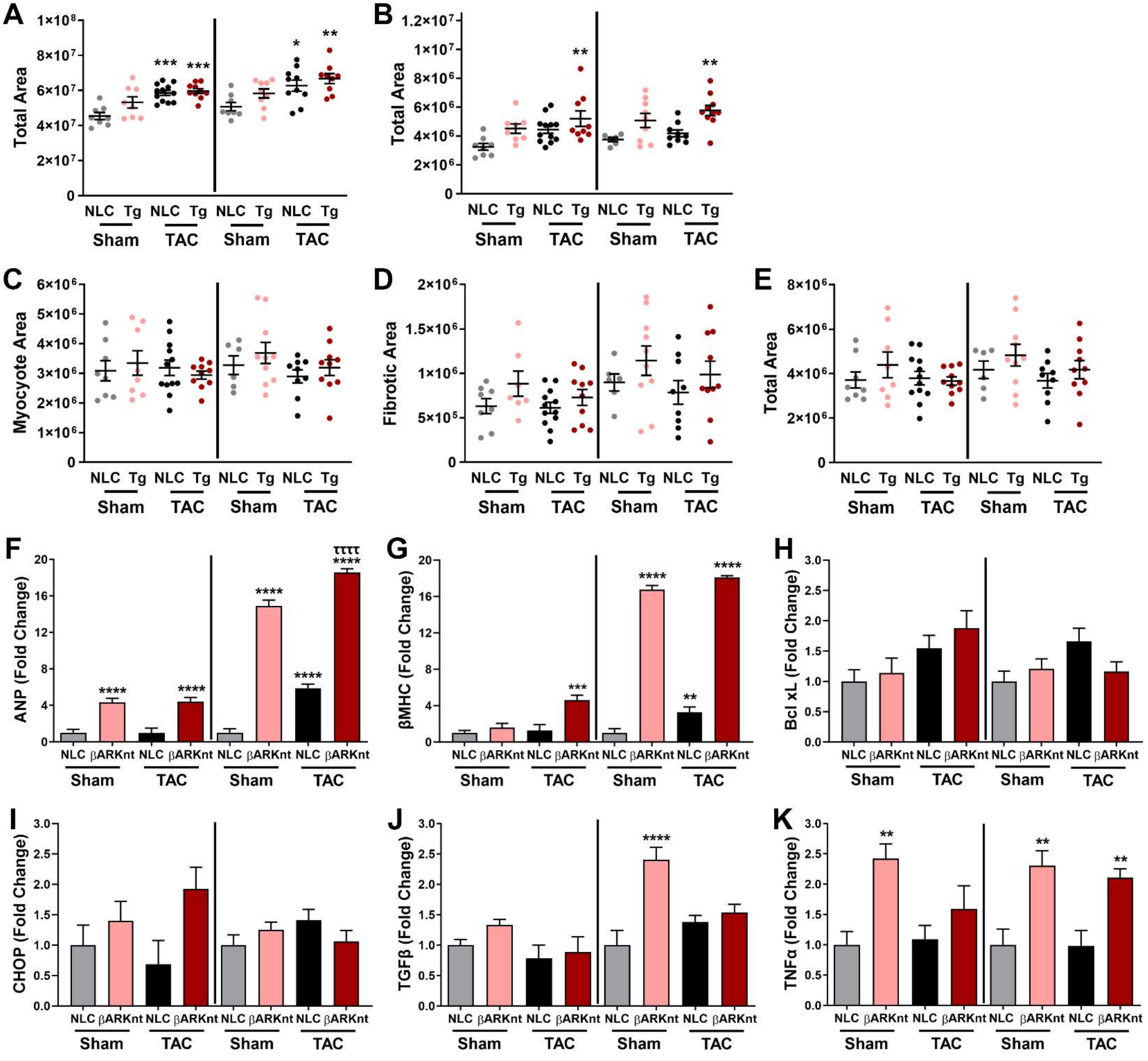


**Figure S3:** *βARKnt hearts show elevated left ventricular and left atrial myocyte and fibrotic area at 4- and 14-weeks after TAC, with equivalently elevated myocyte cross-sectional area but differences in immune cell infiltration.* Quantification of **(A)** total area in the ventricles, **(B)** total area in the left atria, and **(C)** myocyte area, **(D)** fibrotic area, **(E)** total area in the right atria of non-transgenic littermate control (NLC) and TgβARKnt Sham and TAC mice 4 (left) and 14 (right) weeks after surgery. *, $p < 0.05$; **, $p < 0.01$; ***, $p < 0.01$ by one-way ANOVA with Tukey post-hoc test relative to time-matched NLC Sham. n = 9-15 hearts per group. Quantification of RT-PCR data showing fold change in **(F)** atrial natriuretic peptide (ANP), **(G)** β-myosin heavy chain (βMHC), **(H)** B-cell lymphoma-extra large (Bcl-xL), **(I)** C/EBP homologous protein (CHOP), **(J)** transforming growth factor beta (TGFβ), and **(K)** tumor necrosis factor alpha (TNFα) mRNA expression in these mice. **, $p < 0.01$; ***, $p < 0.01$; ****, $p < 0.0001$ by one-way ANOVA with Tukey post-hoc test relative to time-matched NLC Sham. $^{\tau\tau\tau\tau}$, $p < 0.0001$ by one-way ANOVA with Tukey post-hoc test relative to time-matched βARKnt Sham. n = 6-10 hearts per group.

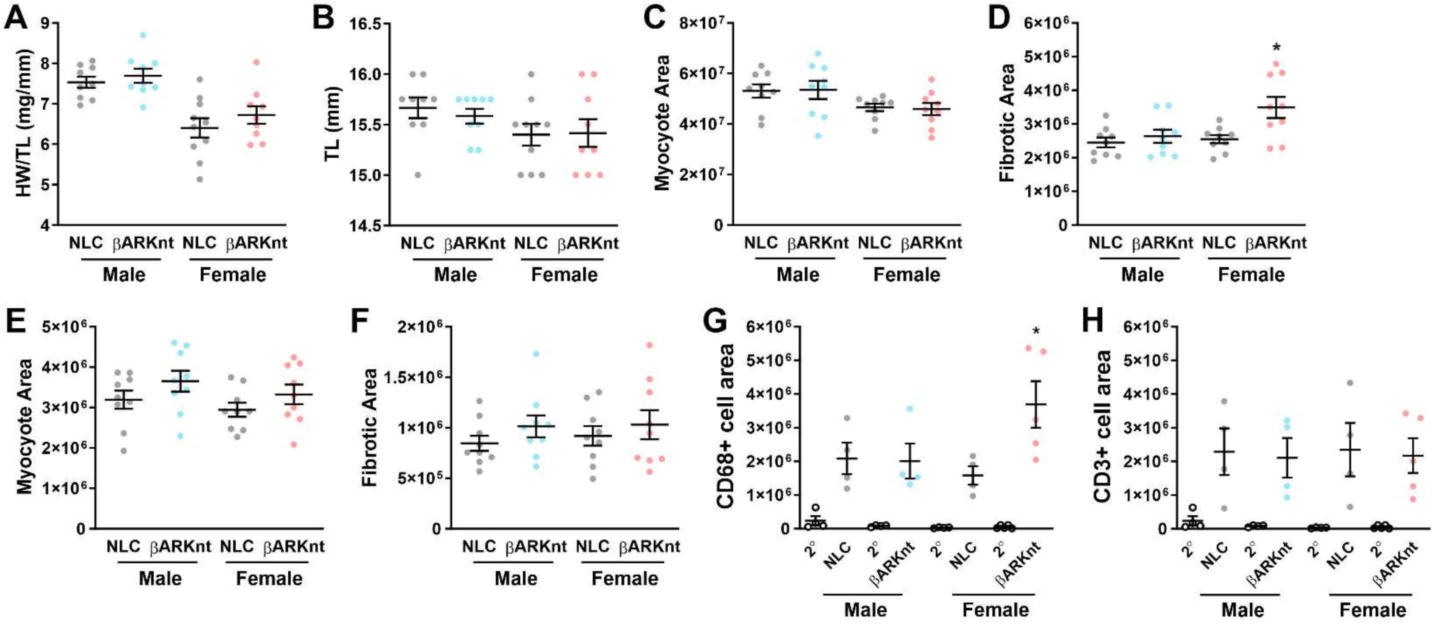


**Figure S4:** *8-week-old βARKnt female mice do not yet exhibit cardiac remodeling but do show increased myocardial fibrosis and macrophage content.* Measures of **(A)** heart weight normalized to tibia length (HW/TL) and **(B)** tibia length in 8-week-old male (left) and female (right) non-transgenic littermate control (NLC) and TgβARKnt mice. n = 9-10 mice per group. Quantification of **(C)** myocyte area and **(D)** fibrotic area in the ventricles of these mice. Quantification of **(E)** myocyte area and **(F)** fibrotic area in the left atria of these mice. *, $p < 0.05$ by one-way ANOVA with Tukey post-hoc test relative to female NLC Sham. n = 9 hearts per group. Quantification of **(G)** CD68 positive cell area and **(H)** CD3 positive cell area in hearts sections from these mice. *, $p < 0.05$ by one-way ANOVA with Tukey post-hoc test relative to female NLC Sham. n = 4-5 hearts per group.

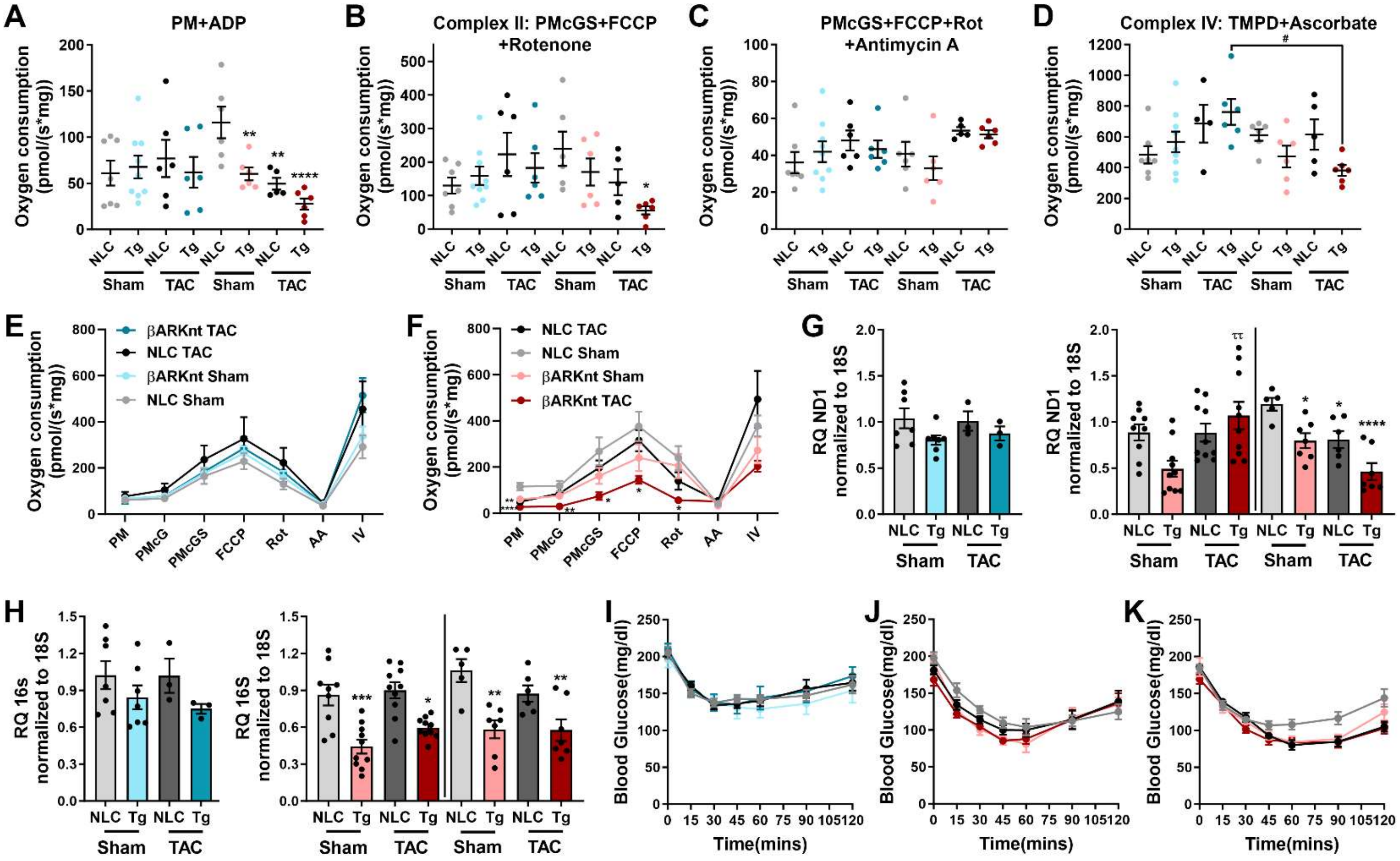


**Figure S5:** *Sex and genotype differences in mitochondrial complex function, resident mitochondrial protein expression, and glucose and insulin handling at 4 and 14 weeks after surgery.* Mitochondrial oxygen consumption in permeabilized cardiac muscle fibers from male and female non-transgenic littermate control (NLC) and TgβARKnt Sham and TAC mice 4 weeks after surgery in response to substrates and inhibitors of components of the electron transport chain (ETC) including **(A)** pyruvate, malate, and ADP (PM+ADP), **(B)** sequential pyruvate, malate, ADP, glutamate, succinate, FCCP, and rotenone (PMcGS+FCCP+Rotenone), **(C)** plus antimycin A, and **(D)** plus subsequent TMPD and ascorbate. Representative tracings of high-resolution respirometry under these conditions in **(E)** male and **(F)** female mice. *, $p < 0.05$; **, $p < 0.01$; ****, $p < 0.0001$ by one-way ANOVA with Tukey post-hoc test relative to sex-matched NLC Sham. $^{\#}$, $p < 0.05$ by one-way ANOVA with Tukey post-hoc test relative to condition-matched male. n = 4-9 hearts per group. Quantification of RT-PCR data showing **(G)** male (left, blue, 4 week) and female (right, red, 4 (left) and 14 (right) week) fold change (RQ = 2−ΔΔCt) in mRNA expression of ND1, and **(H)** male (left, blue, 4 week) and female (right, red, 4 (left) and 14 (right) week) RQ in mRNA expression of 16S in these mice. *, $p < 0.05$; **, $p < 0.01$; ***, $p < 0.01$; ****, $p < 0.0001$ by one-way ANOVA with Tukey post-hoc test relative to time-matched NLC Sham. $^{\tau\tau}$, $p < 0.01$ by one-way ANOVA with Tukey post-hoc test relative to time-matched NLC TAC. n = 3-10 hearts per group. Insulin tolerance test (ITT) measurements of blood glucose levels after IP insulin injection of 0.75U/kg per BW recombinant human insulin in **(I)** male and **(J)** female NLC and βARKnt Sham and TAC mice 4 weeks or **(K)** 14 weeks after surgery. n = 8-13 per group.

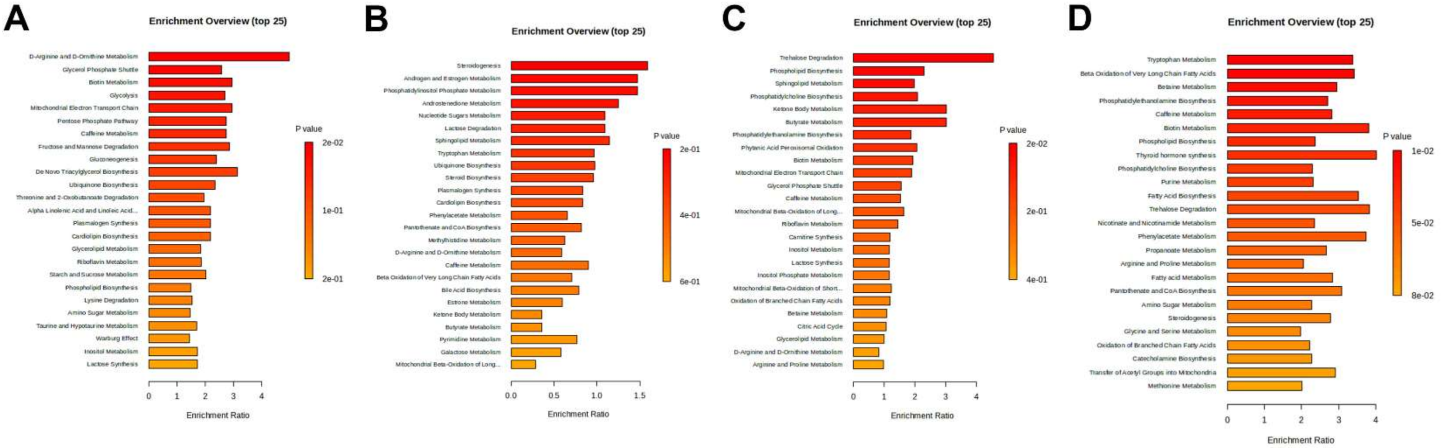


**Figure S6:** *Untargeted serum metabolomics demonstrates strong differences in metabolite prevalence as a factor of genotype and time exposed to pressure overload stress*. Pathway Enrichment analysis results of mapping the full metabolomics dataset from non-transgenic littermate control (NLC) and TgβARKnt serum **(A)** 4 weeks after Sham surgery, **(B)** 4 weeks after TAC surgery, **(C)** 14 weeks after Sham surgery, and **(D)** 14 weeks after TAC surgery with the Small Molecule Pathway Database (SMPDB) using MetaboAnalyst.

**Table S1:** *Untargeted Metabolomics Enrichment Pathway Analysis*

| Groups | Pathway | Total Cmpd | Hits | Statistic Q | FDR |
|---|---|---|---|---|---|
| **βARKnt vs NLC_Sham (4 weeks)** | D-Arginine and D-Ornithine Metabolism | 11 | 1 | 45.121 | 0.56716 |
| | Glycerol Phosphate Shuttle | 11 | 5 | 23.581 | 0.56716 |
| | Biotin Metabolism | 8 | 2 | 26.82 | 0.56716 |
| | Glycolysis | 25 | 4 | 24.552 | 0.56716 |
| | Mitochondrial Electron Transport Chain | 19 | 4 | 26.951 | 0.56716 |
| | Pentose Phosphate Pathway | 29 | 3 | 24.979 | 0.56716 |
| | Caffeine Metabolism | 24 | 4 | 24.908 | 0.56716 |
| | Fructose and Mannose Degradation | 32 | 4 | 25.94 | 0.56716 |
| **βARKnt vs NLC_Sham (14 weeks)** | Trehalose Degradation | 11 | 1 | 41.212 | 0.9797 |
| | Phospholipid Biosynthesis | 29 | 7 | 20.871 | 0.9797 |
| | Sphingolipid Metabolism | 40 | 6 | 17.959 | 0.9797 |
| **βARKnt vs NLC_TAC (14 weeks)** | Tryptophan Metabolism | 60 | 13 | 30.773 | 0.23233 |
| | Beta Oxidation of Very Long Chain Fatty Acids | 17 | 2 | 31.059 | 0.23233 |
| | Betaine Metabolism | 21 | 4 | 26.816 | 0.23233 |
| | Phosphatidylethanolamine Biosynthesis | 12 | 4 | 24.677 | 0.23233 |
| | Caffeine Metabolism | 24 | 4 | 25.627 | 0.23233 |
| | Biotin Metabolism | 8 | 2 | 34.609 | 0.23233 |
| | Phospholipid Biosynthesis | 29 | 7 | 21.582 | 0.23233 |
| | Thyroid hormone synthesis | 13 | 1 | 36.476 | 0.23233 |
| | Phosphatidylcholine Biosynthesis | 14 | 6 | 20.909 | 0.23233 |
| | Purine Metabolism | 74 | 16 | 21.034 | 0.23233 |
| | Fatty Acid Biosynthesis | 35 | 3 | 32.153 | 0.23233 |
| | Trehalose Degradation | 11 | 1 | 34.858 | 0.23233 |
| | Nicotinate and Nicotinamide Metabolism | 37 | 6 | 21.373 | 0.23233 |
| | Phenylacetate Metabolism | 9 | 1 | 33.996 | 0.23233 |

All pathways shown in the table are potential target metabolic pathways with pathway impacts above 0.1. The Holm-Bonferroni method used a Holm p-value of 1 to adjust the FDR with a raw p-value ranging from 0.1461 to 0.049634 of the most significant pathways. Total Cmpd, total number of compounds in the pathway; Hits, the number of matched compounds in the pathway; Statistic Q, the parameter used to adjust the false discovery rate according to the raw p value; FDR, False Discovery Rate adjusted by Holm–Bonferroni method.
